# Natural allelic variations of *Saccharomyces cerevisiae* impact stuck fermentation due to the combined effect of ethanol and temperature; a QTL-mapping study

**DOI:** 10.1101/576835

**Authors:** Philippe Marullo, Pascal Durrens, Emilien Peltier, Margaux Bernard, Chantal Mansour, Denis Dubourdieu^

**Affiliations:** Univ. Bordeaux, ISVV, Unité de recherche OEnologie EA 4577, USC 1366 INRA, Bordeaux INP, 33140 Villenave d’Ornon, France; Biolaffort, 33100 Bordeaux, France; CNRS UMR 5800, Univ. Bordeaux, 33405 Talence, France; Inria Bordeaux Sud-Ouest, joint team Pleiade Inria/INRA/CNRS, 33405 Talence, France

**Keywords:** Key worlds, QTL, *OYE2*, *VHS1*, subtelomeric region, wine yeast, temperature, ethanol

## Abstract

**Background:** Fermentation completion is a major prerequisite in many industrial processes involving the bakery yeast *Saccharomyces cerevisiae.* Stuck fermentations can be due to the combination of many environmental stresses. Among them high temperature and ethanol content are particularly deleterious especially in bioethanol and red wine production. Although the genetic causes of temperature and/or ethanol tolerance were widely investigated in laboratory conditions, few studies investigated natural genetic variations related to stuck fermentations in high gravity matrixes.

**Results:** In this study, three QTLs linked to stuck fermentation in winemaking conditions were identified by using a selective genotyping strategy carried out on a backcrossed population. The precision of mapping allows the identification of two causative genes *VHS1* and *OYE2* characterized by stop-codon insertion. The phenotypic effect of these allelic variations was validated by Reciprocal Hemyzygous Assay in high gravity fermentations (>240 g/L of sugar) carried out at high temperatures (>28°C). Phenotypes impacted were related to the late stage of alcoholic fermentation during the stationary growth phase of yeast.

**Conclusions:** The genes identified are related to molecular functions such as Programed Cell Death, ROS metabolism and respire-fermentative switch and were never related to fermentation efficiency. Our findings open new avenues for better understanding yeast resistance mechanisms involved in high gravity fermentations.

## Background

The yeast *Saccharomyces cerevisiae* presents huge genetic and phenotypic variability that has been recently captured at a large scale level [1]. Beside its worldwide presence in natural habitat, this species is characterized by domesticated strains used in several industrial processes as biofuel, wine, sake, brewery, and bakery [2]. Such strains are specifically adapted to transform sugars in ethanol thought the alcoholic fermentation. One common feature of all industrial strains is the ability to ensure a complete sugar to ethanol conversion since stuck fermentations cause economical prejudice in industry. Most of the environmental factors affecting stuck fermentation have been widely reviewed and partially depend on the industrial application [3][4]. Stuck fermentations may result from the combination of many different stresses including high ethanol content [5, 6], low pH [6, 7], presence of toxins [8, 9], oxygen or nitrogen depletion [10], bacterial contaminations [11, 12], and high temperature [5, 6, 13]. Among others, the combination of high ethanol content and high temperature has been reported to be particularly deleterious for yeast physiology [5, 6, 14]. This is the case for many industrial processes where elevated temperature and high ethanol content are met. Therefore, understanding tolerance mechanisms of fermenting yeast in high temperature and high gravity matrixes is of particular interest. First, in bioethanol industry where Simultaneous Saccharification and Fermentation (SSF) at high temperature (35-41°C) are frequently used [15]. Second, in more traditional food related fermentations; and in particular in red winemaking where the floating cap reaches temperatures significantly higher than those of the bulk liquid, 32–37°C [16, 17].

In order to improve yeast temperature tolerance during alcoholic fermentation, several genetic strategies have been developed such as mutagenesis [18, 19], adaptive evolution [20, 21] and breeding strategies [5, 6] demonstrating that fermentation completion at elevated temperatures is a complex quantitative trait. Beside these applied researches, the ability to growth at high temperature was investigated in laboratory conditions. Particularly tolerant strains were found in clinical samples [22], tropical fruits [23] or *cachaça* brews [24]. These strains, able to growth in laboratory media at up to 42°C, were used for implementing quantitative genetic approaches carried out in standard laboratory media [25]. The genetic basis of High Temperature Growth (HTG) revealed to be particularly complex highlighting the existence of epistatic networks involving multiple genes and their allelic variations [26–29]. Although very efficient, these studies were mostly carried out in physiological conditions that are far from the industrial reality. However, many stresses (including the temperature) impacting the yeast physiology are more stringent during the stationary growth phase since ethanol concentration is higher. In such conditions, the identification of natural genetic variations preventing stuck fermentation were never identified.

In a previous work, we constructed by successive backcrosses a Nearly Isogenic Lineage (NIL) improved for its fermentation performance at 28°C [5]. In this lineage, nearly 93% of the genome is identical to one parental strain showing stuck fermentation at elevated temperature. The remaining 7% of the genome contains heterozygous genetic regions that prevent stuck fermentation. In the present work, this genetic material was used for carrying out a QTL mapping using a selective genotyping strategy. Three main QTL were identified and two of them were dissected at the gene level leading to the identification of two causative genes encoding the proteins Oye2p and Vhs1p. The third locus mapped was the subtelomeric region of the chromosome XV that could play a role in this complex trait.

## Results

### Genetic material and experimental design

Among many others, the temperature is an impacting factor that influences the fermentation completion [30]. In a previous study, we demonstrated that this parameter induced stuck fermentations for many wine industrial starters when they are steadily fermented at 28°C. In contrast, in the same media, most of them achieved the fermentation when the temperature was maintained at 24°C. For another group of strains, the temperature change did not affect the fermentation completion. These observations suggested a differential susceptibility to temperature in high gravity medium that was defined as thermo-sensitive/tolerant trait [5]. Among various wine yeast strains, this phenotypic discrepancy is particularly high for the meiotic segregants B-1A and G-4A, which are derived from commercial starters Actiflore BO213 and Zymaflore F10, respectively (Laffort, FRANCE) (Table 1). In a breeding program, the hybrid H4 was obtained by successive backcrosses using the tolerant strain, B-1A as the donor and the sensitive strain, G-4A as the recipient strain (see Figure 1A). These backcrosses were driven by selecting recursively the meiotic segregants showing the best fermentation performance at 28°C [5]. The resulting hybrid H4 had a strong genetic similarity (~93%) with the recipient background G-4A but also inherited some genetic regions from B-1A conferring a more efficient fermentation at 28°C (Figure 1A).

**Table 1.**
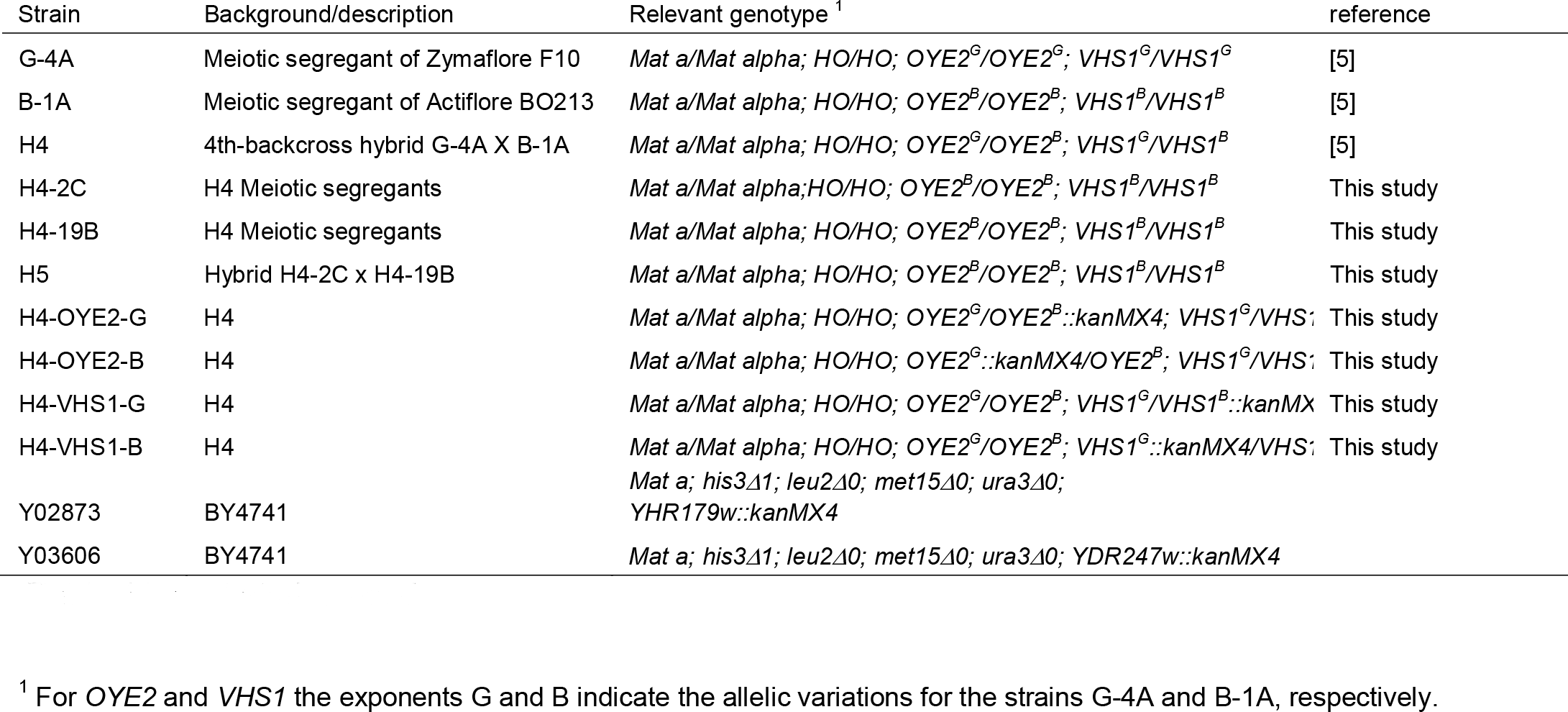
Yeast strains used

**Figure 1.**
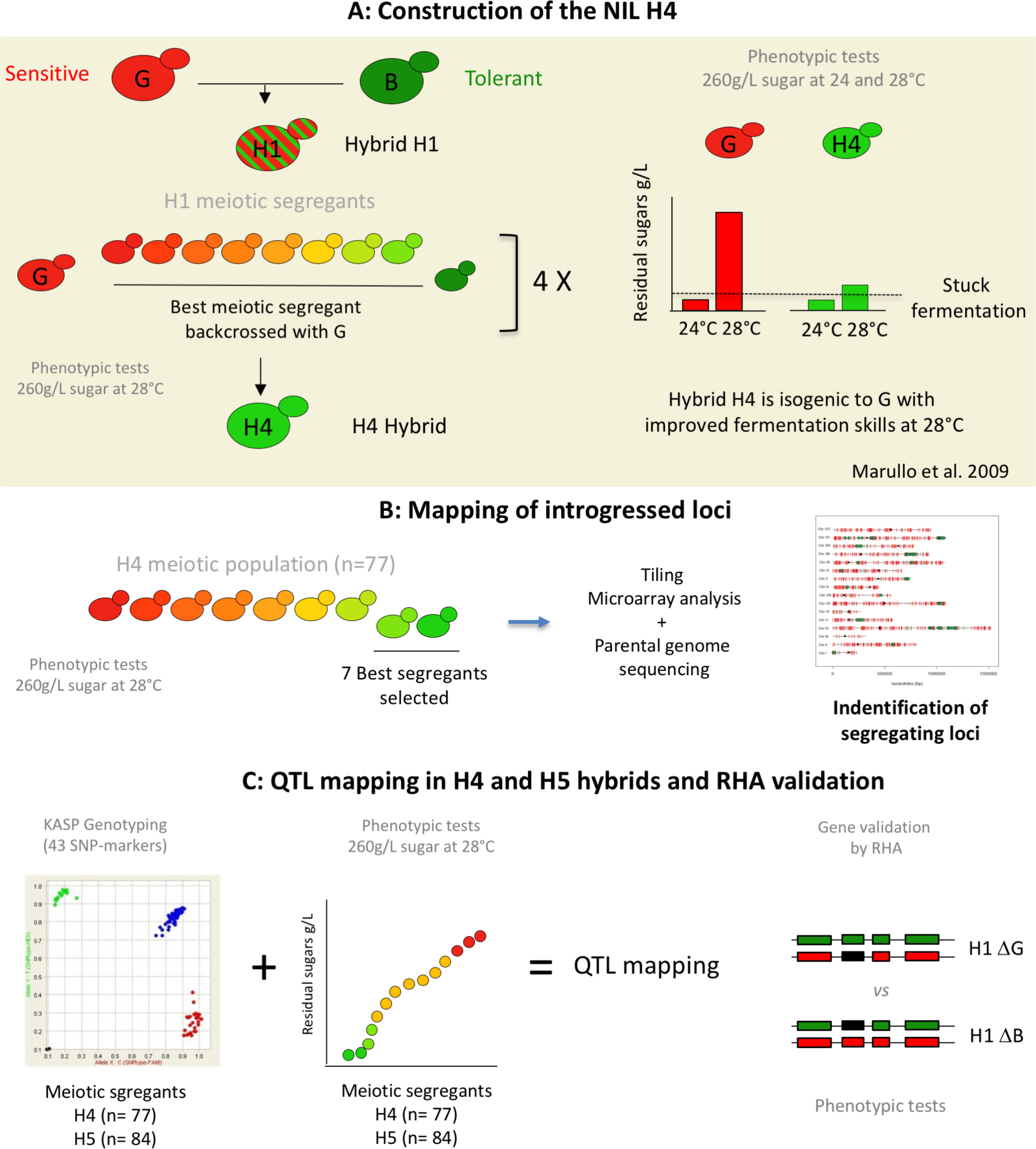
Genetic material and experimental design. The panel A summarizes the construction of the genetic material used in this study. The H4 hybrid was obtained by a backcross program using the parental strains G-4A (G) and B-1A (B). The F1-hybrid was sporulated and the resulting segregants were phenotyped for their fermentation performance at 28°C. The segregant leaving the smallest quantity of residual sugars was cross with the strain G-4A. This procedure was recurrently done four time in order to get the hybrid H4 that constitutes the starting point of this present study [5]. Phenotypic comparison of the hybrid H4 and G illustrates that fermentation efficiency of H4 was specifically improved at 28°C as reported by Marullo *et al.* [5]. The panel B describes the strategy used for mapping the chromosomal portion of the strain B-1A present in the hybrid H4. In order to narrow the most relevant regions, a selective genotyping approach was applied. 77 H4-segregants were fermented and the seven best ones were genotyped by combining Tiling Microarray (Affymetrix®) and whole genome sequencing. The panel C describes the QTL mapping strategy applied that was carried out by developing qPCR-based markers (KASP^TM^ technology) in order to achieve a linkage analysis using up to 160 segregants. Candidates genes identified were then validated by reciprocal hemizygosity assay (RHA)

The aim of the present study is to identify the genetic determinisms explaining the phenotypic variance observed in this nearly isogenic population by applying QTL mapping approach. The overall strategy is presented in the Figure 1 (B and C). Initially, the phenotypic segregation of fermentation traits was investigated in 77-segregants of H4. Then, seven extreme individuals leaving the lowest concentration of residual sugars were individually genotyped by Affymetrix^®^ Tiling microarray. This selective genotyping step allowed the localization of genomic regions inherited from B-1A that have been introgressed in the G-4A genome during the backcross. Finally, numerous segregants (~160) belonging to two backcrossed hybrids (H4 and H5) were genotyped using Kompetitive Allele Specific PCR markers (KASP^TM^). A linkage analysis identified three QTLs, two them were molecularly dissected by Reciprocal Hemizygous Assay.

### Phenotypic characterization of H4 progeny

The parental strains (B-1A, G-4A), the hybrid H4, and 77 H4-meiotic segregants were fermented in a synthetic grape must containing 260g/L of sugar at 28°C (see methods). Most of the strains showed stuck fermentation due to the harsh conditions applied. The overall phenotypic characterization was carried out by measuring eight quantitative traits (Table 2). According to the phenotype, the heritability *h^2^* in the H4 progeny ranged from 2.5 to 86.9 %. Kinetic traits in relation with the early part of alcoholic fermentation (*LP*, *T35*, *T50*) were poorly heritable and are not statistically different within the parental strains. None of these traits were further investigated due to their low heritability. The lack of segregation within the offspring suggests that all the segregants share a similar phenotype for the first part of the fermentation which correspond to the growth phase. This observation has been previously reported for one particularly tolerant segregants of H4 showing growth parameters very similar to the parental strain G-4A [5]. In contrast, traits linked to the late part of the fermentation (*T70*, *rate 50-70*, *ethanol produced*, *CO_2_max*, *Residual Sugars (RS*)) had a high variability. This is the case of the Residual Sugars at the end of the alcoholic fermentation (Figure 2A). For this trait, the parental strains values are 0.1 and 30.3 g/L for B-1A and G-4A, respectively. A complete overview of the trait segregation is given for all the trait investigated (Additional file 1 and 2). The contrasted segregation between early and late fermentation traits indicates that the underlining genetic determinisms would be more linked to physiological mechanisms related to the stationary growth phase. The QTL mapping was only applied to two phenotypes (Residual Sugar and T70) since most of them are strongly correlated (Additional file 3).

**Table 2.**
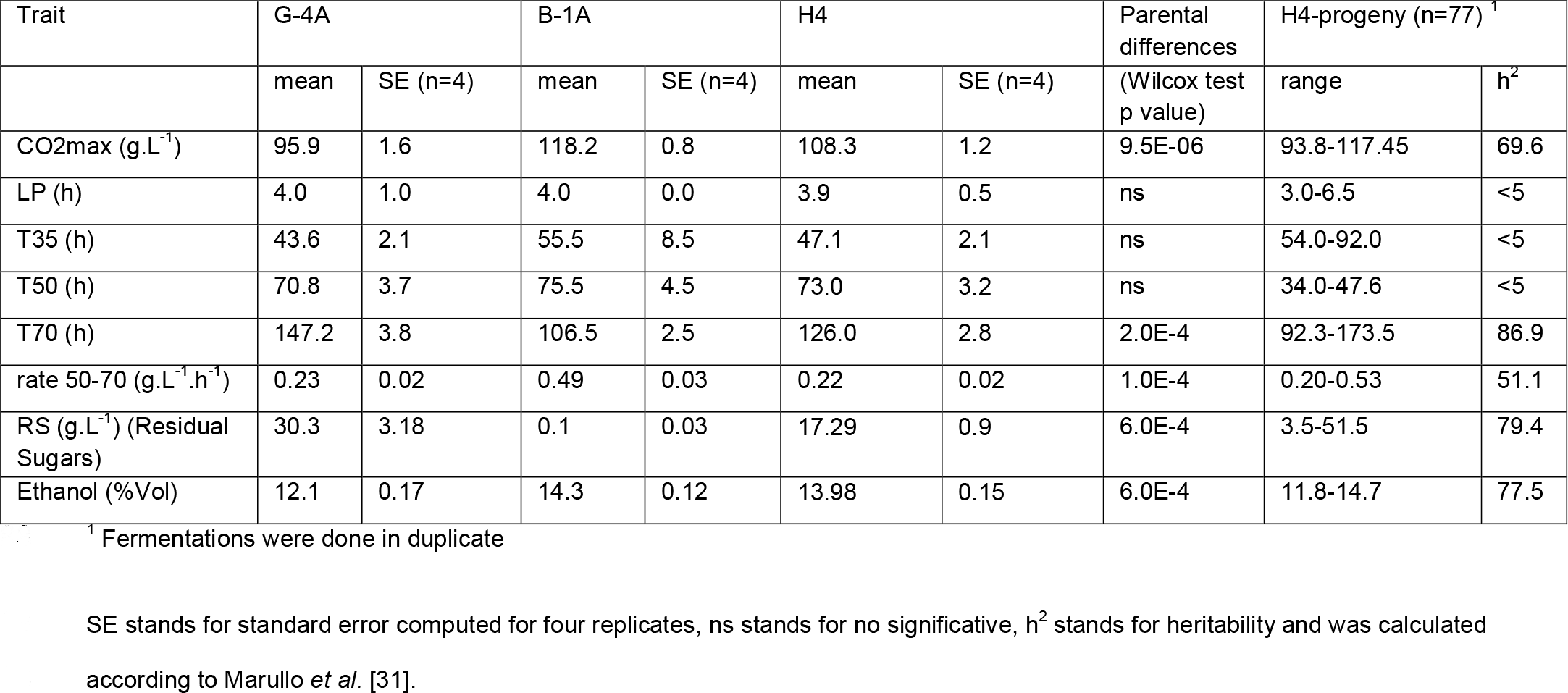
Phenotypes of parental strains and for the H4 progeny.

**Figure 2.**
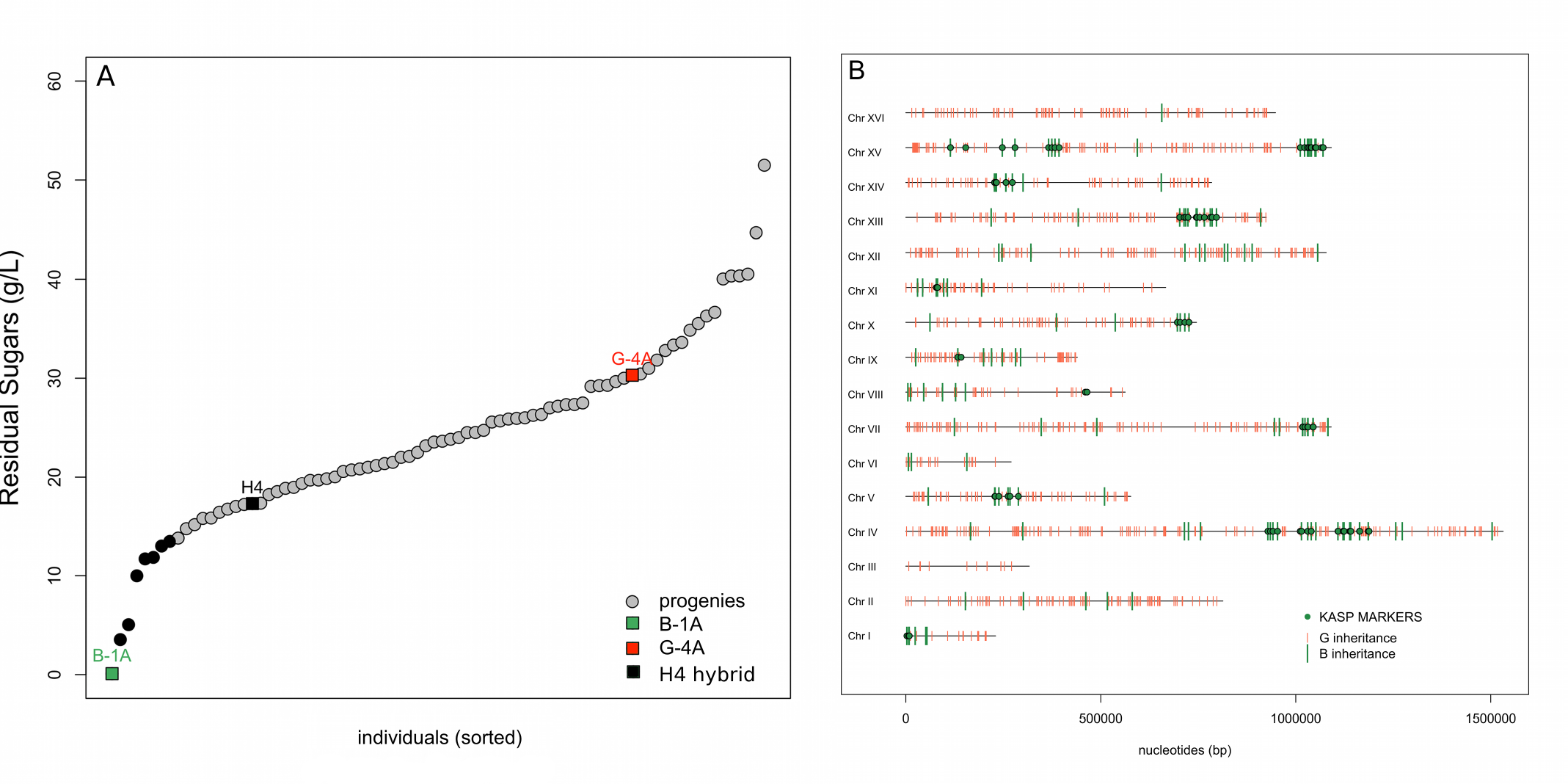
QTL regions narrowed by selective genotyping. Panel A. Distribution of the residual sugars found at the end of the alcoholic fermentation for the 77 H4-segregants and for the parental strains. The average values of parental strains and H4-hybrid were indicated by green (B-1A), red (G-4A) and black squares (H4-hybrid). The segregants values are the means of experimental duplicates, the seven best progenies (black dots) were selected for narrowing the QTL regions. Panel B, Physical map of all the B-1A and G-4A specific markers inherited in the seven H4 progenies. Each thick is one of the 1204 bi-allelic markers selected. The B and G alleles are shown in green and red, respectively. The green dots are the SNP that were found in more than four segregants defining 12 chromosomal regions.

### Narrowing introgressed loci by selective genotyping with Affymetrix^®^ Tiling microarray

In order to identify QTLs explaining the phenotypic variation observed, a selective genotyping approach was implemented. First, the genomic DNA of the parental strains G-4A and B-1A were hybridized on Yeast Tiling Microarray (YTM). Using the algorithm SNP Scanner described by Gresham et al. [32]; 18601 and 12848 SNP were detected with respect to the reference genome (Saccharomyces cerevisiae S288C strain, R49.1.1, 2005) for the strains B-1A and G-4A, respectively. Among these SNP, 3397 non-common positions were found defining putative markers between the parental strains (Additional File 4). The correct assignation of these predicted SNP was verified by checking their position with the complete sequence of the parental strains obtained by whole genome sequencing taking as reference the (Saccharomyces cerevisiae S288C strain, (version Apr2011/sacCer3) (Additional File 4). As the algorithm was not able to predict exactly the position of the SNP, a search window was defined with various intervals ranging from 5 to 20 bp.

More than 80 % of the detected SNP were located at least than 10 bases of the position predicted by YTM. However, only 1204 predicted SNP were correctly assigned meaning that in our experiment the False Discovery Rate of YTM was close to 65%. Nevertheless, the 1204 validated SNP constitutes reliable bi-allelic markers covering the most part of the genome. According to the inheritance of the parental strains B-1A and G-4A, these markers were thereafter named “B” and “G”, respectively. The inheritance of this set of markers was investigated in the H4 segregants. In order to reduce the genotyping cost, only the seven H4 segregants were individually genotyped by YTM. These segregants were selected on the basis of their ability to achieved the most part of the alcoholic fermentation according to their RS values (Figure 2A). This number represent the best decile of the H4 progeny which is sufficient to narrow the main genetic regions containing QTLs [33]. Due to the recurrent backcrosses operated, only 192 markers (green ticks) inherited from B-1A were detected in the genome of the seven progenies genotyped. The remaining 1012-markers were inherited from the parental strain G-4A (red ticks). The B-specific markers were mainly clustered in 12 genomic regions localized in 11 chromosomes (Figure 2B). Half of them (89 green dots) were found in more than 4 of the 7 progenies genotyped. Since they are more frequently found in the best progenies, those regions are supposed to encompass the B-specific markers conferring a more complete fermentation at 28°C. According to the segregant, the proportion of B-markers was very similar, ranging between 14.3 and 16.9 %. This ratio is a bit higher than expected for a 4 times backcrossed hybrid but clearly confirms that the genetic imprinting of parent B-1A has been reduced by the backcross procedure as previously demonstrated by a microsatellite analysis [5]. From the 192 B-markers identified, we selected a subset of markers in order to genotype a larger population. On the basis of parental genome sequence, 43 KASP^TM^ markers localized in the 12 genomic regions were designed (Figure 2B); their genomic positions are given in Additional file 5.

### Sequential QTL mapping in two NIL populations identifies three loci linked to stuck fermentation

The 77 segregants of the backcross hybrid H4 were genotyped by using the KASP^TM^ markers (LGC genomic company, UK). This technique allows detecting SNP inheritance by using a qPCR method with labeled primers [34]. The correct Mendelian segregation of 43 SNP in this population was confirmed (chi^2^ test, α=0.05) as well as the homozygous status of each segregant (>99 % of the SNP). A linkage analysis was carried out by using a non-parametric test (Wilcoxon test, α=0.05) with a significant threshold fixed by 1000 permutations as previously described [35]. The use of non-parametric test was justified by the heterogeneity of variance of the phenotype investigated. Two main QTLs localized on the chromosome IV and VIII were mapped for phenotypes *RS* and *T70* (Figure 3 A and B). The maximum linkage values were found for the markers IV_953 and VIII_464. For both loci, the B-1A inheritance conferred an improved phenotype, which is in accordance with parental strains phenotypes (Figure 3 C and D). The part of variance explained by those QTLs ranged between 15.6 and 25.8 % according to the trait and the locus (Table 3). The analysis of variance of the linear model described an additive effect without interaction.

**Table 3.**
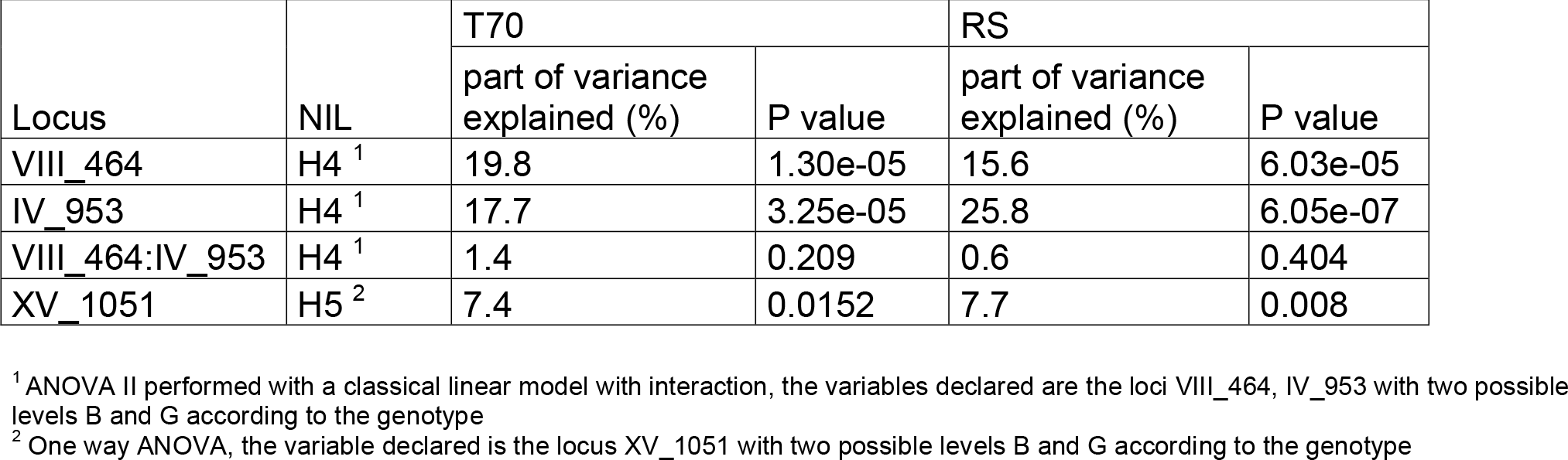
QTL effects and part of variance explained.

**Figure 3.**
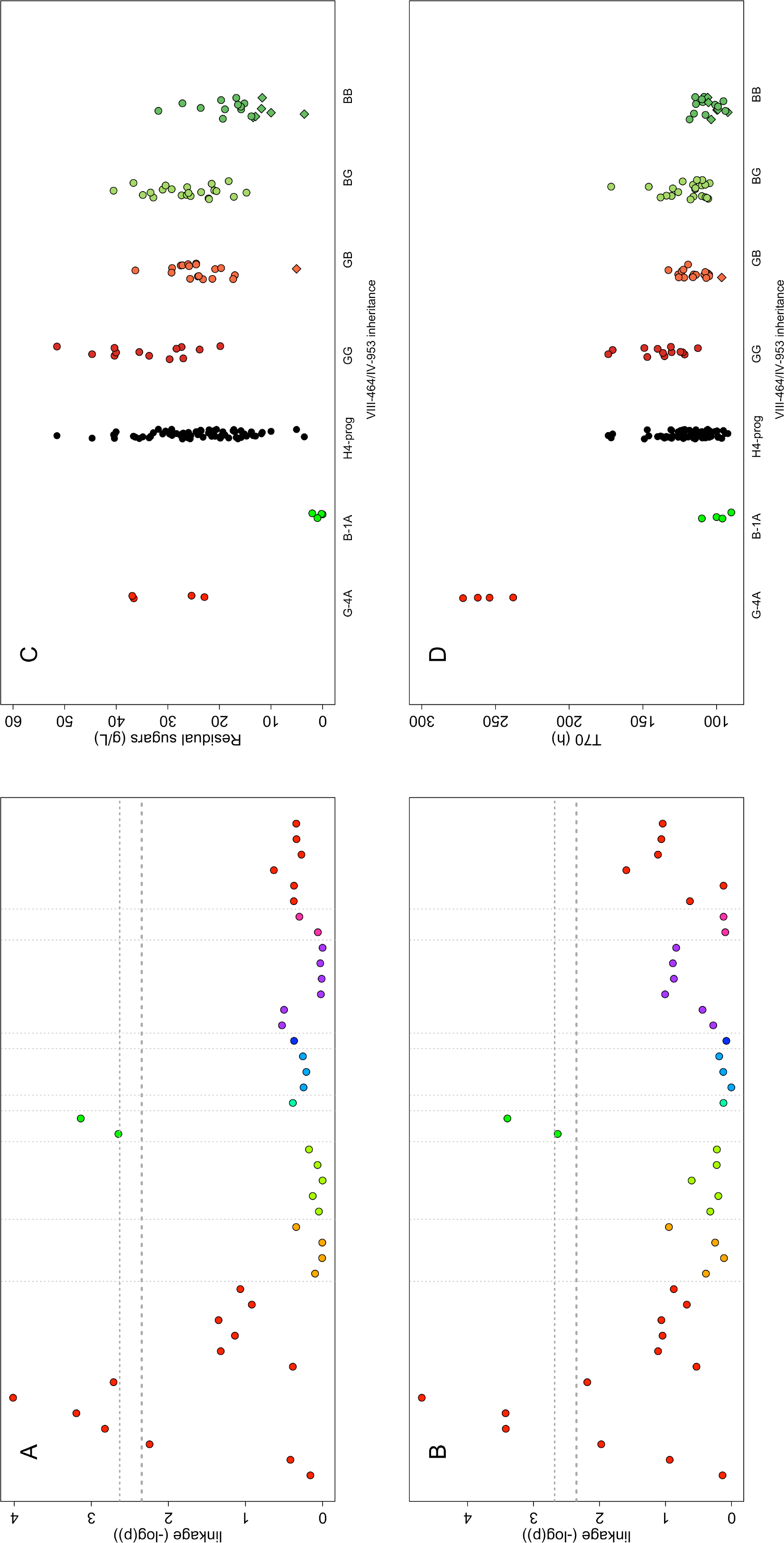
Linkage analysis in the H4 progeny. Panel A and B show the linkage score expressed in – log of pvalue (Wilcox-Mann-Withney test) for the 43 qPCR markers used for QTL mapping of Residual sugars and T70, respectively. The dot colors represent markers on different chromosomes. The p-value thresholds of False discovery Rate (FDR 10% and 5%) were estimated by permutation tests (n=1000) and are shown by tight and wide dotted lines, respectively. Panel C and B. Trait distribution among the H4 progeny according to the inheritance at the loci VII-464 and IV-953 for Residual Sugars (g/L) and T70 (h), respectively. The parental values are indicated at the left part of the dot plot. The seven progenies selected were indicated by diamonds symbols. The letters G and B stands for G-4A and B-1A inheritance, respectively.

This first genetic mapping captures only 40% of the total variance observed within H4 progeny suggesting that other QTLs playing a minor role were not yet detected. More complex mapping methods integrating the QTL position as cofactors failed to detect any other loci (data not shown), likely due to the relative small number of segregants analyzed and the low density of the map. According to the strategy proposed by Sinha *et al.* [27], the effect of the two major QTLs was removed by performing an additional cross. We selected two H4 segregants (H4-19B and H4-2C) showing a B-alleles inheritance for the QTLs IV_953 and VIII_464. These clones were selected in order to maximize their phenotypic distance for *RS* (close to 30 g/L). The resulting hybrid H5 was heterozygous for only 23 loci localized in 8 chromosomal regions (Additional Table 5). A population of 84 segregants of the H5 hybrid was then isolated, phenotyped and genotyped in the same way than for H4 segregants. The phenotypic segregation of this population is given in the Table 4. Although the trait heritability was lower than for H4 progeny, some traits of interest such *RS* and *T70* are clearly inheritable and showed a wide segregation. This lower heritability is likely due to the fact that traits were measured without replicates in order to maximize the number of segregants tested. This choice can be justify by the fact the most important factor affecting QTL-mapping efficiency is the number of individuals rather than the phenotype measurment accuracy [33]. A new linkage analysis allowed the detection of one additional QTL localized in the subtelomeric region of chromosome XV (Figure 4 A). The maximum peak linkage was found for the marker XV_1051. Surprisingly, for this locus the G-4A allele was linked to a more efficient fermentation for both *RS* and *T70* traits. One-way analysis of variance indicates that only 7.5% of the total variance was explained by this QTL in the H5 progeny (Table 3).

**Table 4.**
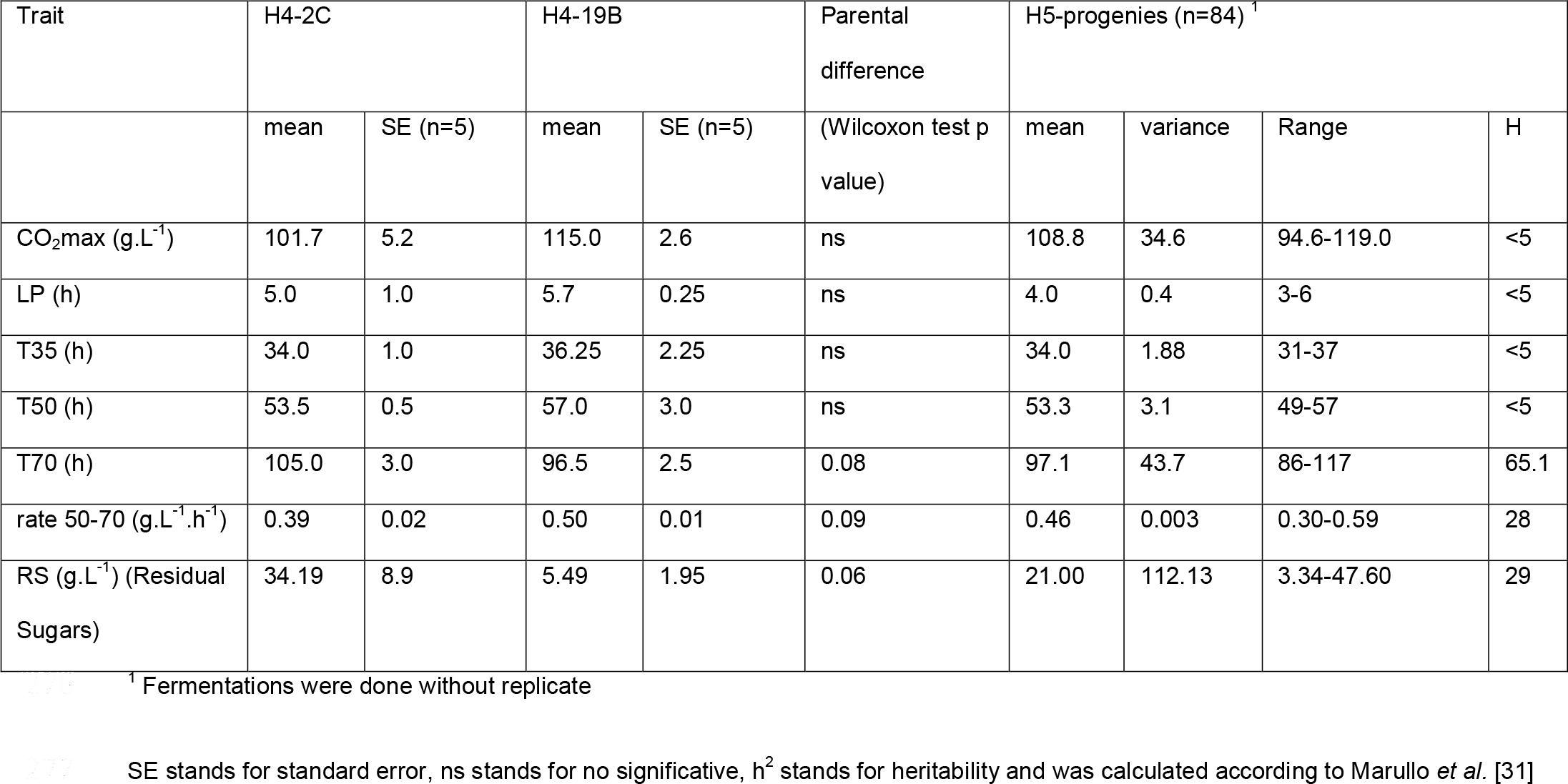
H5 Phenotypes

**Table 5.**
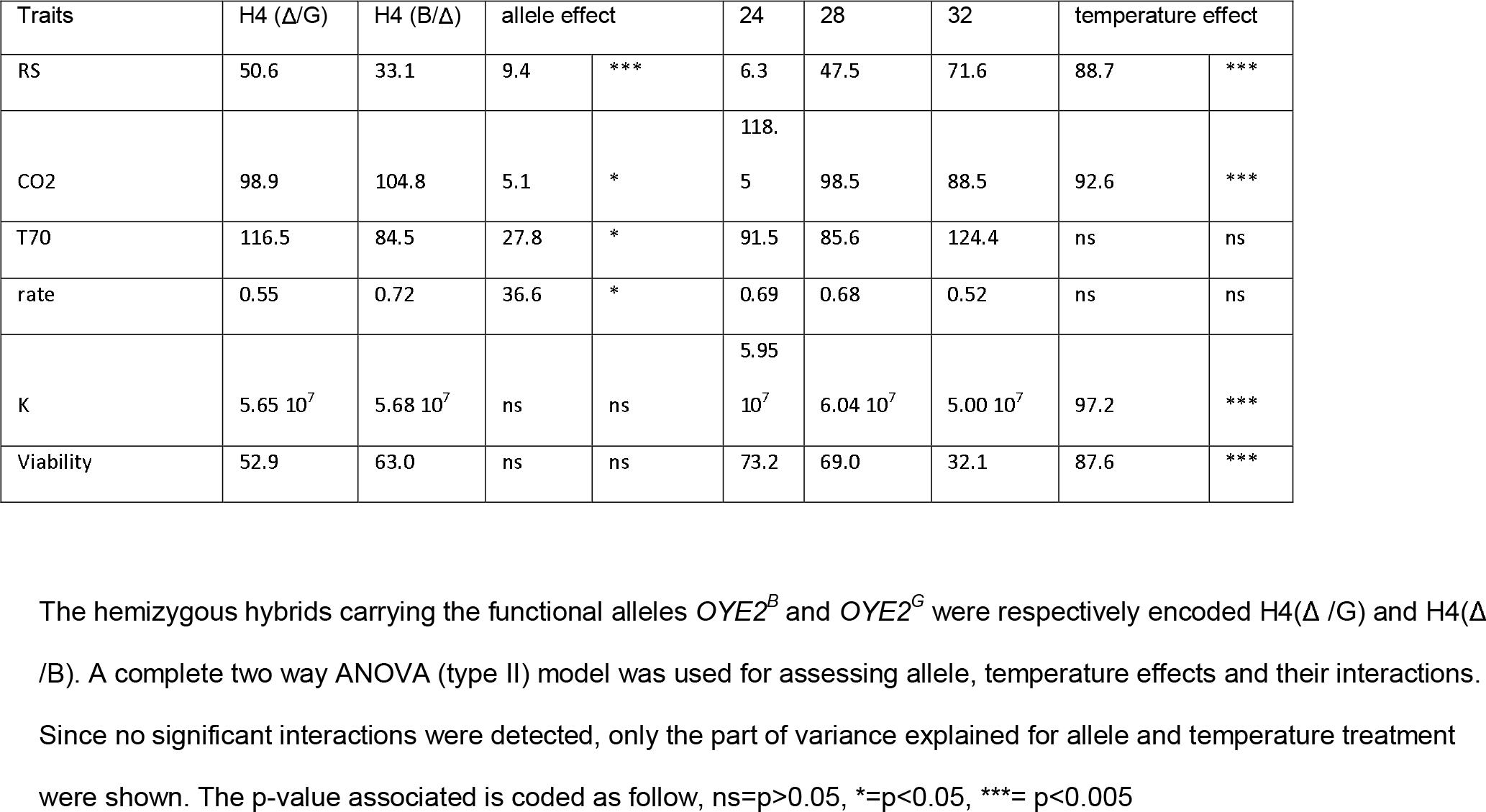
Effect of temperature and *OYE2* alleles on the main fermentation parameters

**Figure 4.**
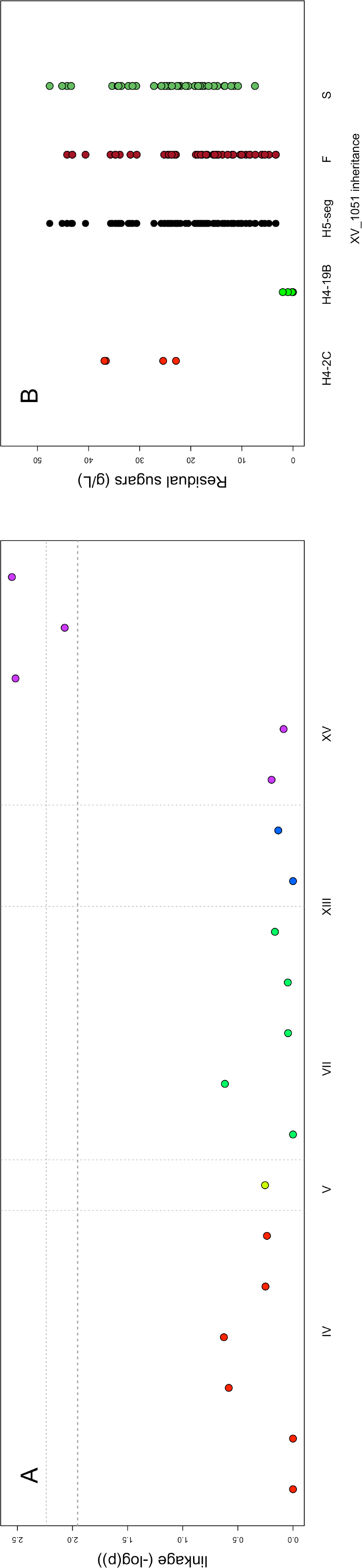
Mapping of the minor QTL XV in the H5 progeny. Panel A shows the linkage score expressed in – log of p-value (Wilcox-Mann-Withney test) for the 19 qPCR markers used for QTL mapping of Residual sugars in the H5 progeny. The dot colors represent markers on different chromosomes. The p-value threshold of False discovery Rate (FDR 10% and 5%) was estimated by permutation tests (n=1000) and are shown out by tight and wide dotted lines, respectively. Panel B. Residual Sugars (g/L) distribution among the H5 progeny according to the inheritance at the loci XV_1051. The parental values (H4-2C) and H4-19B) are indicated at the left part of the dot plot. The letters G and B stands for G-4A and B-1A inheritance, respectively.

After this additional linkage analysis, three QTLs (VIII_464, IV_953 and XV_1051) explaining nearly 50 % of the phenotypic variation have been significantly detected.

### Impact of the NADPH oxidoreductase Oye2p on stuck fermentation in high sugars and temperatures conditions

We first investigated the QTL VIII_464 by analyzing the genomic sequence of both parental strains neighboring 15 kb from the best marker found. This region (456000 to 472000 bp) encompassed 7 genes; four of them (*STB5*, *OYE2*, *YHR180W*, *YHR182W*) showed non-synonymous SNP between the parental strains (Additional file 6). At less than 2 kb of the QTL peak, a deletion at the position genomic position 462732 (c.229_230delTC) produced a frame-shift mutation in the *OYE2* gene of the parental strain G-4A (p.Ser77fsTer95). The resulting ORF produces a truncated protein of only 95 amino acids instead of the 400 expected in the full-length protein. This two-bases deletion was thereafter named *OYE2^G^* allele. In contrast, the strain B-1A has the same sequence as the reference genome (S288c) producing a full-length protein form (thereafter named *OYE2^B^*). By screening genome databases, we did not detect this specific deletion in other 100 strains (data not shown). However, two other strains carry missense polymorphisms that generate truncated Oye2p proteins OS104 (p.Gly73fs) and S294 (p.Gln176*) (Figure 5A).

**Figure 5.**
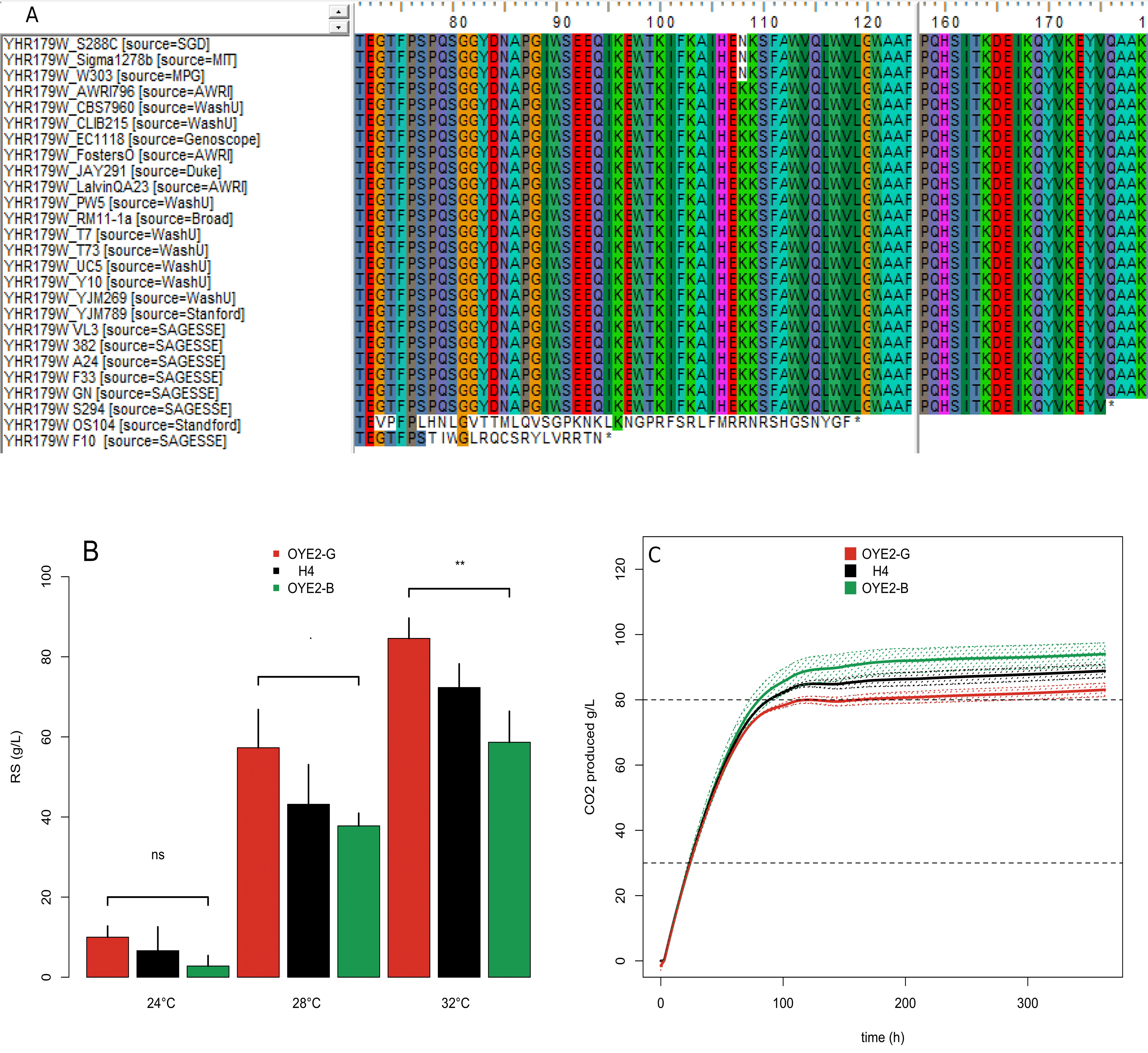
Physiological effect of the *OYE2* alleles. Panel A. Sequence alignments of *Saccharomyces cerevisiae Oye2p* proteins. The strains F10 (parental strain of G-4A), OS104 and S294 show stop-codon insertion at different positions. Panel B. The bar plots represent the average values of residual sugars after isothermal fermentations carried out at 24, 30 and 32°C in both hemyzygous hybrids and the native H4 hybrid. The genotypes Δ*OYE2^B^*::*KanMx4/OYE2^G^*, *OYE2*^B^/Δ*OYE2^G^::KanMx4,* and *OYE2*^B^/*OYE2^G^* are shown in red, green and black, respectively. Bars represent standard error of five repetitions, the statistical differences between the hemizygous was tested by a Wilcoxon-Mann-Whitney Test (the p value is coded as follow, ‘ns’=p>0.05, ‘.’=p<0.1, ‘**’=p<0.01. Panel C. Fermentation kinetics (CO_2_ produced time course) for the same strains and with the same color key.

To test the impact of this candidate gene, Reciprocal Hemizygous Assay (RHA) [25] was implemented. This method allows the comparison of each parental allele in the H4-hybrid background. The strains H4-OYE2-G and H4-OYE2-B were obtained by using a deletion cassette. These hemizygous hybrids had genotypes Δ*OYE2^B^*::*KanMx4/OYE2^G^* and *OYE2*^B^/Δ*OYE2^G^::KanMx4,* respectively (Table 1). Their fermentation performances were compared at different fermentation temperatures (24, 28 and 32°C). In addition to the CO_2_ fermentation kinetics, biomass samples were taken in order to estimate the maximal population reached as well as the cell viability at 70% of the fermentation. An analysis of variance (type II) reveals that both temperature and *OYE2*-allele nature impacted many phenotypes. The temperature effect accounts for the major part of the phenotypic variance confirming its deleterious effect on the fermentation completion in high gravity conditions.

Beside this notorious environmental effect, our results demonstrated that the nature of the *OYE2* allele significantly affected the fermentation kinetics (*T70* and rate), residual sugar content (*RS)* and the total amount of CO_2_ produced (Figure 5B and C). In contrast, neither growth, biomass content, nor cellular viability were different between the hemizygous hybrids (Additional file 7). Therefore, the physiological impact of *Oye2p* concerns more fermentation activity than the cell growth or viability. In standard laboratory conditions, the *OYE2* hemizygous showed exactly the same fitness (data not shown). For the *RS* measured at 28°C, the phenotypic difference between H4-OYE2-G and H4-OYE2-B hybrids was close to 16 g/L. By splitting the H4-progenies in two groups according to their inheritance for *OYE2* marker, the average phenotypic discrepancy within the two groups is only 9.2 g/L. Epistatic relationships within other genes might explain why the *OYE2* effect in H4-offspring is lower than that observed in the hemizygous hybrids. This finding suggests that other genes close to *OYE2* might control this phenotype. Nevertheless, our data suggest that the gene *OYE2* strongly contributes to the QTL’s effect.

As the mutation (p.Ser77fsTer95) is the unique *Oye2p* peptidic variation found between the parental strains, the OYE2^G^ allele (p.Ser77fsTer95) should be responsible of the deleterious effect observed. In the hybrid H4, phenotypes observed are quite similar to those observed in the hemizygous hybrid H4-OYE2-B suggesting the recessive nature of this mutation (Figure 5B and C). In other to evaluate more generally the implication of the *OYE2* gene in wine fermentations, we assayed in laboratory strain background (BY4741) the physiological impact of its full deletion using the strain Δ*oye2* (Y02873). The fermentation conditions were adapted by reducing the initial sugar content (180g/ instead of 260g/L) since the laboratory strain is much less adapted than industrial backgrounds (Additional File 8). The *OYE2* deletion impacted both kinetics and RS content at 24° and 32°C confirming the importance of this gene in the fermentation resistance.

### Impact of the protein kinase Vhs1p on stuck fermentation in high sugars and temperatures conditions

The second QTL localized in the right arm of chromosome IV was also dissected. During the backcross procedure, three distinct introgressed regions were inherited form B-1A, encompassing a very large portion of chromosome IV (Figure 2B). Only the central zone was statistically linked to the phenotype with a maximum peak detected for the marker IV_953. In this genomic region (945500 to 957800), nine non-synonymous variations were found within the parental strains affecting 7 genes (Additional file 6). The most striking mutation was a nucleotide substitution C to A at the position g.957128 producing a stop-codon (p.Tyr372*) in the gene *VHS1*. This gene encodes for a cytoplasmic serine/threonine protein kinase. Interestingly the stop-codon came from the parental strain B-1A conferring a more efficient fermentation. The resulting protein is truncated for 93 C-terminal amino acids but conserves its catalytic domain. This allelic variation was not detected in any other strains (n=100). Thereafter this mutation is named *VHS1^B^* in opposition to the wild type allele *VHS1^G^* carried by the parental strain G-4A and the reference strain S288c. The effect of this gene was validated by RHA by constructing the hemizygous hybrids H4-VHS1-G and H4-VHS1-B (Table 1). In the experimental conditions used for QTL mapping (260 g/L of sugar, 28°C), we did not observe a significant effect of this gene due the high variance observed within repetitions. However, by reducing the sugar concentration to 240 g/L and increasing the fermentation temperature up to 32°C a significant effect of the *VHS1^B^* allele was observed Figure 6 A and B. As for *OYE2,* no differences are found neither for growth, biomass production, nor viability whatever the culture medium used (synthetic grape juice or laboratory medium). The weak effect observed is likely due to the fact that other genes in this genetic region also impact this phenotype. The hybrid H4 has the same phenotypic level than the hemizygous hybrid H4-VHS1-G suggesting that the beneficial allele *VHS1^B^* is mostly recessive. Alike for *OYE2,* we verified the *VHS1* effect in another genetic background (BY4741) applying milder fermentation conditions (180g/L of sugar). The deletion mutant Δ*vhs1* (Y03606) showed a significative reduction of fermentation efficiency respect to the control by leaving more residual sugars (Figure 6 C) and having a slower fermentation kinetics (Figure 6 D). In contrast to the results observed in the H4 background, the deletion effect of *VHS1* was not observed at 28°C but at 24°C. This could be explained by the very sluggish fermentation kinetics of laboratory strain that was unable to consume 40% of the total sugar. These additional results demonstrated that a loss of function of this protein is deleterious for fermentation efficiency. Strikingly, the effect of total deletion of *VHS1* in BY background contrasted with the partial C-terminal deletion observed in H4. Indeed, the *VHS1^B^* allele has a positive effect on the fermentation efficiency. Altogether, reciprocal hemizygous and functional genetic analyses of *VHS1* provide new insights of this poorly characterized kinase. Interestingly a positive natural allelic variation associated to fermentation resistance in high temperature and ethanol conditions has been identified. The weak effect of this mutation suggested that *VHS1* is not the unique gene explaining the effect of this QTL. Therefore, other allelic variations physically linked to VHS1 are likely involved and remain to be identified.

**Figure 6.**
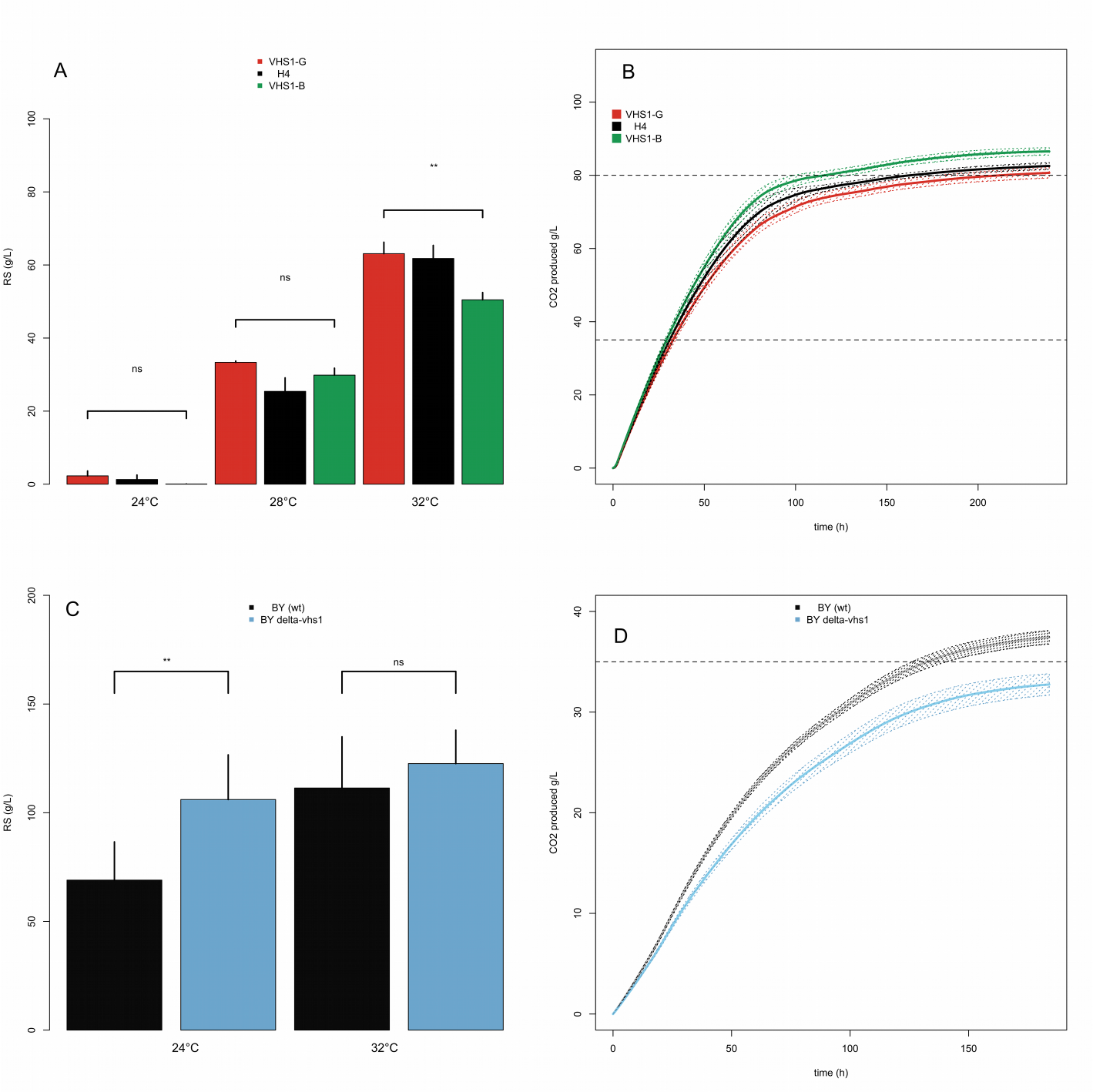
Physiological effect of *VHS1* alleles. Panel A. The bar plots represent the average values of residual sugars after isothermal fermentations carried out at 24, 30 and 32°C in both hemyzygous hybrids and the native H4 hybrid. The genotypes Δ*VHS1^B^*::*KanMx4/VHS1^G^*, *VHS1*^B^/Δ*VHS1^G^::KanMx4,* and *VHS1*^B^/*VHS1^G^* are shownin red, green an black, respectively. Bars represent standard error of five repetitions, the statistical differences between the hemizygous was tested by a Wilcoxon-Mann-Whitney Test (the p value is coded as follow, ‘ns’=p>0.05, ‘.’=p<0.1, ‘**’=p<0.01. Panel B. Fermentation kinetics (CO_2_ produced time course) for the same strains and with the same color key. Panel C The bar plots represent the average values of residual sugars after isothermal fermentations carried out at 24, and 32°C in the laboratory strain background (BY4741). The genotypes Δ*vhs1* and *VHS1(wt)* were shown in blue and grey, respectively. Bars represent standard error of five repetitions, the statistical differences between the strains was tested by a Wilcoxon-Mann-Whitney Test (the p value is coded as follow, ‘ns’=p>0.05, ‘**’=p<0.01). Panel D. Fermentation kinetics (CO_2_ produced time course at 24°C) for the same strains and with the same color key.

## Discussion

### Efficient mapping of three QTLs linked to stuck fermentation triggered by high ethanol and temperature conditions

In this work, we investigated the genetic causes preventing stuck fermentations in high sugars and temperatures conditions by using a QTL mapping analysis. In order to reduce the genotyping cost, the QTL mapping was carried out on a Nearly Isogenic Lineage (NIL). This lineage is derived from meiotic segregants of the hybrid H4, which has been obtained by successive backcrosses, using as parental donor the strain B-1A, and as recipient one the strain G-4A. In order to identify the main regions inherited from B-1A, Yeast Tiling Microarrays (YTM) were used for genotyping the seven best segregants of H4. This selective genotyping strategy is routinely used in quantitative genetics and allows the reduction of genotyping effort by increasing the detection power of major QTLs [33]. Selective genotyping could be achieved by Bulk Segregant Analysis which requires pooling large set of extreme segregants [36]. In yeast, this strategy has been particularly useful for mapping QTLs linked to fitness differences by allowing an easy selection of numerous individuals with extreme phenotypes [37]. In our experimental conditions, all the progenies showed a similar growth/viability and are mostly different for their fermentation kinetics and their residual sugars values. Therefore, an easy selection based on fitness or viability was not possible and all of them were phenotyped individually. The selection of only seven segregants representing the tail of RS distribution (the best 10%) was sufficient to map 12 genomic regions by using Tiling microarray. Once the main introgressed regions have been localized, we applied an additional filter using the parental genome sequences. This filter was necessary since the SNPs detected by Tiling microarray were not all reliable (FDR= 65%). A subset of 43 bi-allelic markers segregating in a Mendelian way across the H4 population was successfully defined. Those markers were selected in genomic regions mostly inherited from B-1A strain assuming the fact that most of tolerance alleles should be brought by this parental strain. By genotyping 77 segregants, 2 main QTLs (VIII_464 and IV_953) were detected confirming the efficiency of our strategy. Despite the low-density map used, it is noteworthy that both QTL mapped are very close to the causative genes identified (*OYE2* and *VHS1*) less than 4 kb.

However, this first linkage analysis captured only 40 % of the total phenotypic variation observed. The identification of only two major QTLs explaining 40% of the total variance observed in the H4-progeny is consistent with quantitative genetics theory. In fact, most of the QTL-studies fail to capture all the genetic variability especially due to the strong epistatic relations within QTLs [38]. In order to find out other QTLs, the effect of those major loci was removed by achieving another cross among phenotypically distant progenies having the same inheritance (B) for the markers VII_464 and IV_953. This additional analysis leads to map a third locus localized at the end of chromosome XV (peak linkage at the marker XV_1051). Due to the weak effect of this QTL, we did not investigate the causative genes of this region. However, some details recently reported stroke our attention.

The end of chromosome XV was linked with the genomic region C acquired by horizontal transfer which is present by most of the wine yeast [39]. Recently, it was demonstrated that the genes *FOT1-2* of this region derived from *Torulaspora microellipsoides* confers evolutionary advantages in grape must fermentations [40]. However, both parental strains have a full copy of the gene *FOT1-2* suggesting that the cause of their phenotypic discrepancy is elsewhere. By comparing the read coverage of the parental strains, we found that B-1A has a 12-kb deletion in the right subtelomeric part of the chromosome XV (Additional file 9). This deletion encompassed the genes *FIT2, FIT3, FRE5, YOR385W, PHR1,* and *YOR387W* and has been previously described for other yeast strains [41]. Since the allele of the strain B-1A confers temperature sensitivity, the lack of one or many genes of this region could be directly involved. Similarly, a recent work linked the cold temperature tolerance with various subtelomeric regions of chromosome XIII, XV, and XVI having such kind of deletion [42]. Our results suggest that the right arm of chromosome XV might play a similar role for fermentation efficiency. However, further genetic analyses are obviously necessary for confirming this hypothesis and identifying the causative gene(s) involved. Some segregants carrying all the three positive alleles constitute a relevant genetic material for carrying out breeding programs in order to improve the performance of other strains by using marker assisted selection strategies [43]

### New insights in the physiological role of the old yellow reductase Oye2p

The first QTL mapped (VIII_464) has a major impact on stuck fermentation since it explains more that 25% of the total variance observed for residual sugar (Table 4). By using a Reciprocal Hemizygosity Analysis, the role of the gene *OYE2* (*YHL179c*) was clearly established. The allele *OYE2^G^* strongly impaired the success of alcoholic fermentation in anaerobic conditions since the parental strain G-1A has a two nucleotides deletion (c.229_230delTC) which generates a truncated protein of 94 amino acids (instead of the 400 expected). In the G-1A background, this allele conferred the same phenotype than the null mutation *oye2::KanMx4* (data not shown). The phenotypic effect of *OYE2* gene was also confirmed in another unrelated background (S288c) demonstrating its relevant impact on the alcoholic fermentation resistance (Additional File 8). In the industrial background, the *OYE2^G^* effect is only observed during the alcoholic fermentation of high quantity of sugars when the temperature exceeds 32°C (Figure 5). The phenotype discrepancy appeared after 67 h of fermentation when more than 70 % of sugar has been consumed. Indeed, no significant difference in fermentation kinetics were observed during the first part of the alcoholic fermentation. This result indicates that the physiological role of this protein can be observed only in drastic environmental conditions (32°C, 260 g/L of sugar). This finding highlights the relevance of achieving applied genetic studies in conditions that match as much as possible with industrial practices.

The protein Oye2p is a conserved NADPH oxydoreductase [44] belonging to the large family of flavoenzymes that has a growing interest in biocatalysis [45]. Despite several studies, the physiological role of Oye2p remains unclear. This could be due to the phenotypic conditions used that were mostly carried out in exponential phase conditions (*e.g.* YPD at ~28°C). Large-scale functional genomics suggested that Oye2p should have a possible role in cytoskeleton assembly [46] as well as in oxidative stress response [47, 48]. This mitochondrion-associated protein was also characterized for its anti-apoptotic effect by lowering Programed Cell Death (PCD) after various oxidative treatments [48]. Interestingly, a connection between oxidative stress conditions and heat shock has been previously established for this gene [49]. The effect of *oye2* deletion on cell viability after a transient heat shock (50°C, 20 min) was only observed when the cells are sampled during the stationary phase but not during exponential growth [49]. This result is consistent with the effect detected here that occurred only at a relatively high temperature (32°C).

Our findings suggest that *OYE2* could play a key role in the mechanisms of cell resistance in the late part of alcoholic fermentation. Those mechanisms could be linked with the protective effect of Oye2p against ROS (Reactive Oxygen Species). Indeed, it has been demonstrated that in late steps of wine alcoholic fermentation, important amounts of ROS are observed [50]. Since high temperature [51] and ethanol [52] promote ROS production in *S. cerevisiae*, the harsh conditions met in our experiment may have emphasized this phenomenon. The direct role of ROS accumulation for triggering the cell death in yeast is still debated [53]. However, cellular mechanisms preventing the increase of ROS are supposed to delay the induction of PCD during the alcoholic fermentation. The protein Oye2p could play this protective role by reducing the ROS production and their damages in anaerobic conditions. Connecting Programmed Cell Death and fermentation performance is a promising route for better understanding stuck fermentations in industry [54]. Recently, it has been shown that the deletion of the pro apoptotic protein Sch9p as well as the inhibition of TOR complex prevent cell death in micronutrient starvations [55]. The potential role of *OYE2* to prevent PCD in oxidative conditions, led us to measure the cell viability as well as the biomass production of hemizygous hybrids during the alcoholic fermentation. However, no significant differences were detected (Additional file 7, Table 6) indicating that the allelic forms tested did not affect cell viability in our conditions. This could be explained by the relative low amount of nitrogen used (190 mg.NL^-1^), which contrasted with the high levels used by Duc *et al.* [55]. Although the cell death seems not directly triggered in our conditions, the absence of functional Oye2p could have other deleterious consequences since this protein has been described to have a role in the regulation of actin polymerization [56]; the modulation of cellular glutathione content [48], and the sterol metabolism [57]. Finally from an enological point of view, the presence of non-functional Oye2p could also impact the volatile composition of wine since this protein has been described to reduce geraniol into citronellol [58].

### Evidence of a truncated form of the protein Vhs1p impacting fermentation efficiency

We dissected at the gene level a second QTL (IV_953) also linked to the ability to achieve the later part of the fermentation. Reciprocal hemizygous analysis demonstrates the physiological effect of the gene *VHS1* (Figure 6) and partially validates at the gene level the impact of this QTL. Compare to the locus VIII_464, this genomic region is quite large (> 400 kb) (Table 3) and four markers spaced by 25 kb were positively linked to the phenotype. Previous studies in *S. cerevisiae* [7, 25] already described these particularly large QTLs, which could be due to the presence of several causative genes brought by both parental strains. This could be the case here since other genes having a close phenotypic relation with temperature (*HSP48*) or acid pH (*RKR4*) resistances are located in the same region. Therefore, *VHS1* is likely not the unique gene of this QTL that impact the phenotypes investigated. By comparing the genomic sequence of parental strains, we detected close to the QTL peak (IV_953), a non-sense mutation (c.1116C>A) in the coding sequence of *VHS1* of the strain B1-A. This gene encodes a cytoplasmic serine/threonine protein kinase.

Strikingly, the favorable allele (*VHS1^B^*) generates a truncated C-terminal protein of 371 amino acid instead of 461. The 90 amino acids deleted do not encompass the protein kinase domain (PS50011) suggesting that the truncated form might conserve its serine-threonine kinase activity. This effect is recessive as indicated by the RHA assay. The positive effect of the truncated allele of *VHS1^B^* contrasted with the full deletion of *VHS1* done in another background (Figure 6 C and D). Indeed, in the laboratory background the full gene deletion has a deleterious impact on fermentation fitness. The role of temperature in the expressivity of this gene remains unclear. Although the Repriprocal hemizygous analysis indicates an effect only at 32°C (Figure 6A and B), the *VHS1* deletion is deleterious in S288C background whatever the temperature. The molecular function of this kinase has been partially characterized in a recent study [59]. This protein is involved in the regulation of the Snf1p, a central regulator of the carbon metabolism. By phosphorylating the protein *Sip5p,* Vhs1p indirectly activates the regulator Snf1p. Therefore, Vhs1p indirectly promotes the fermentation metabolism and might contribute to maintain the expression level of fermentative genes during the second part of the alcoholic fermentation. Further analyses are required for deciphering the precise physiological role of this kinase and in particular the role of the *VHS1^B^* allele that confers a fermentation resistance in high sugar and temperature conditions.

## Conclusion

In this study we identified by a QTL mapping approach, two natural allelic variations impacting the fermentation performance of industrial yeast in sugar rich media (>250g /L of glucose-fructose). A third locus encompassing the deletion of 6 subtelomeric genes has been also detected. The combination of selective genotyping and the further selection of few markers did not impact the precision of QTL mapping leading to capture nearly 50 % of the total phenotypic variation.

Interestingly, the genes identified are related to molecular functions such as programed cell death, ROS metabolism, and respire-fermentative switch. These findings open new avenues for studying the mechanisms of fermentation resistance and especially those caused by a high temperature and a high sugar content.

## Material and methods

### Yeast strains and culture conditions

All the *Saccharomyces cerevisiae* strains used during this work were listed in Table 1. All strains were propagated at 30°C on YPD medium (1% yeast extract, 1% peptone, 2% glucose) solidified with 2% agar when required. When necessary the antibiotic G418 (Sigma-Aldrich, St Louis, Missouri, USA) was added at a final concentration of 100 μg/ml. The construction of the backcross hybrid H4 and the description of the initial parental strains G-4A and B-1A have been previously reported [5]. The hybrid H5 was obtained by crossing two selected meiotic segregants of H4 (H4-2C and H4-19B) on the basis of their genotype as detailed in result section.

A collection of 77 and 84 meiotic segregants respectively derived from H4 and H5 was obtained by spore dissection using a Singer manual apparatus. The segregants obtained are diploid fully homozygous cells due to the homothallic character of the hybrids H4 and H5 (*HO*/*HO*). Due to their nearly isogenic nature, the germination rate of the hybrids H4 and H5 were close to 100% and all the segregants showed the same fitness on laboratory media.

### Phenotype measurement

#### Alcoholic fermentation

Fermentations were run in the KP medium, a synthetic grape juice which mimics a standard grape juice [60]. This medium was sterilized by filtration through a 0.45 μm nitrate-cellulose membrane (Millipore, Molsheim, France). Phenotypes were measured in a medium containing 260 g.L^-1^ of sugars (50% fructose-50% glucose) with a diluted amount of anaerobic growth factors (S+A-conditions) as described previously [5]. For the fermentation with the BY strains, the amount of sugar was reduced to 160 g/L. moreover in such experiment the strain auxotrophies were complemented by adding uracil (20 mg/L), methionine (20 mg/L), leucine (30 mg/L), histidine (30 mg/L) in the synthetic media. A solution stock of [ergosterol (15 µg.L^-1^), sodium oleate (5 µg/L) and 1mL Tween 80/ethanol (1:1, v/v)] was added in the medium with a 5000 fold dilution. Pre-cultures were run in Erlenmeyer flasks for 24h at 24°C under orbital agitation (150 rpm) in the fermentation media filtered and diluted 1:1 with milli-Q water. The inoculum concentration was 10^6^ viable cells per ml. Fermentations were then run in closed 150mL glass-reactors, locked to maintain anaerobiosis, with permanent stirring (300 rpm) at 28°C. The CO_2_ released was monitored by measurement of glass-reactor weight loss regularly and expressed in g.L^-1^. Raw weight lost data were smoothed by a *Loess* function allowing the estimation of various kinetics parameters. The *CO2max* (g.L^-1^) was the maximal amount of CO_2_ produced during the fermentation, the *LP* (h) was the lag phase time before the CO_2_ production rate was higher than 0.05 g.L^-1^.h^-1^, the *T35*, *T50*, *T70* are the time necessary to reach 35, 50 and 70% of the maximal amount of CO_2_ expected (125 g.L^-1^); the *rate 50-70* (g.L^-1^.h^-1^) was the rate of CO2 released between *T70* and *T50*. Due to the important number of progenies analyzed, two distinct batches of fermentation were carried out. To estimate eventual block effects parental strains (G-4A, B-1A for H4) and (H4-2C, H4-19B for H5) were fermented in both batches two (or three) times leading to get 4 (or 5) replicates of each parent and their relative hybrids. No significative block effects were observed for the traits analyzed (data not shown). For the H4 segregants, all the fermentations were done in duplicate, for the H5 segregants only one series was carried out.

#### Fermentation analyses

At the end of the alcoholic fermentation, the synthetic wines were analyzed for basic enological parameters. *Glucose* and *Fructose* consumed were estimated by enzymatic assay using the kit n° 10139106035 according to manufacturer protocol (R-Biopharm, Germany) and the *RS* (Residual Sugars) was computed as the sum of remaining glucose and fructose (expressed in g.L^-1^). The linearity of this method was presented in the Additional file 10 and was recently published [61]. *Ethanol* (%Vol) was measured by infrared reflectance using an Infra-Analyzer 450 (Technicon, Plaisir, France).

### Whole genome sequencing of parental strains

Whole genome sequences of strains B-1A and G-4A were obtained by using Illumina pair-end sequencing. Briefly, genomic DNA was extracted from a saturated culture of 100 ml under anaerobic condition (YPD) using the genomic tip-100 kit (Qiagen, Courtaboeuf, FRANCE). Paired-end Illumina sequencing libraries were prepared from sonicated genomic DNA according to manufacturer protocols (Genomic DNA Sample Preparation) and were realized by the Genomic and Transcriptomic Facility of Bordeaux, FRANCE. Sequencing was performed on Illumina Genome Analyzer IIx (Illumina, Palo Alto, CA) with a read length of 54 bp. Raw reads data have been deposited in the SRA at NCBI with the accession number PRJNA419624. The genome coverage was respectively 45X and 34X for B-1A and G-4A, respectively. After reads quality trimming and filtration step, each strain was aligned to the reference genome of *Saccharomyces cerevisiae* S288c (version Apr2011/sacCer3) using “Bowtie2” with default parameters. Single Nucleotide Polymorphisms (SNPs) were called using Samtools *mpileup* with *mapping quality* ≥30, *base quality* ≥20, and *varFilte*r depth ≥10. Single amino-acid polymorphisms were identified using *snpEff* [62] requiring quality QUAL ≥30 and genotype GEN[*] GQ ≥ 20. Using this procedure, we defined a set of 9829 high-quality SNP (Q>30, homozygous) named WGS-SNP and given in Additional File 4.

### Selective genotyping using Yeast Tiling Microarrays (YTM)

The genomic DNA of the diploid parental strains (G-4A, B-1A) and of seven H4 segregants was isolated as previously described [31] and hybridized onto *GeneChip S. cerevisiae* Tiling Array 1.0 from Affymetrix (Palo Alto, CA) following the protocol detailed by Gresham *et al.* [32]. Hybridization and microarray scanning were performed by the ProfileXpert platform (Lyon, France). For each parental strain, two independent hybridizations were carried out. Single Nucleotide Polymorphism (SNP) and short Insertion Deletion (INDEL) were searched using the *SNPscanner* program [32] built for scanning SNP on the reference genome release R49-1-1 (2005). In order to reduce the heterogeneity of fluorescence signal between each microarray, the Z score of hybridization signal was calculated according to [63]. The prediction threshold of z scored-transformed data was higher than 2.5 and only regions extending for at least 10 nucleotides above the signal threshold were conserved. This approach allows recovering 3354 putative markers within the strains. Using the same procedure, seven segregants of the hybrid H4 were genotyped. The compiled set of YTM markers inherited from B-1A and G-4A found in the seven segregants was listed in the Additional File 3. The *Perl* and *R* scripts used for computing these dataset are available on request.

### PCR-based KASP™ genotyping of H4 and H5 progenies

Genomic DNA of segregants were extracted using the Genome Wizard (Promega, France) kit using the modified conditions described by Zimmer et al. [35]. The inheritance of 43 SNP was investigated using the KASP genotyping assay based on the use of one universal FRET cassette reporter system. Primers design and genotyping were performed by LGC genomics (Hertz, UK).

### Reciprocal hemizygosity assay

Gene deletion were carried out by homologous recombination using deletion cassette constructed by PCR using as template the genomic DNA of Euroscarf collection strains (Euroscarf, Franckfurt, Germany). The *OYE2* and *VHS1* deletion cassette were obtained using as template the genomic DNA of the strains Y02873 and Y03606, respectively. All the constructs were verified by both insertion and deletion PCR test. The insertion test consists to positively amplify by PCR a fragment containing the 5′ part of the KanMx4 cassette and ~600 bp of the 5′-flanking region of the target gene. Deletion test consists in the absence of amplification of a central portion of the target gene. The hybrid H4 and the parental strains B-1A and G-4A were transformed using the lithium acetate protocol described by Gietz and Schiestl [49]. The allele identity in hemizygous hybrids was tested by sequencing. All the primers used are listed in Additionnal File 11. For each hemizygous hybrid assay at least two independent clones of each genotype were tested.

### Determination of cell viability and concentration

The cell concentration (cells/ml) as well the cell viability were estimated at 70% of the total CO_2_ expected using a flow cytometer (Quanta SC MPL, Beckman Coulter, Fullerton, California), equipped with a 488nm laser (22mW) and a 670 nm long-pass filter. Samples were diluted in McIlvaine buffer pH=4.0 (0.1M citric acid, 0.2M sodium phosphate dibasic) added with propidium iodide (0.3% v/v) in order to stain dead cells (FL3 channel).

### Graphical and statistical analyses

All the statistical and graphical analyses were carried out using the R program (R version 3.3.3 2017-03-06). The custom R scripts used are available on request. The global heritability of each trait (h^2^) was estimated as described previously [60]. The correlation among traits were estimated using a Pearson test corrected by a Bonferroni’s test using the R package *Coortest*. Linkage analysis was carried out according to the non-parametric method (Wilcoxon-Mann-Whitney) used by Zimmer *et al.* [35]; and by calculating for each trait a significant threshold by 1000 permutation tests (α=0.05). The QTLs, genes, and temperature effects were estimated by standard complete linear models (with interaction) and further analyzed by ANOVA (type II). For each variable, the homogeneity of the variance was assessed using a Levene test (*car* package) and the normality of residual distribution was controlled using a Shapiro test (α>0.01). Duncan’s multiple comparison was used to determine which means differ significantly (Duncan’s multiple comparison, α=0.05) using the *agricolae* package. When required, pairwise comparisons were carried out using the Wilcoxon test with at least four independent repetitions.

## Additional files

### Additional File 1

Fermentation kinetics of the 77 segregants and the two parental strains G-4A (red) and B-1A (green).

### Additional File 2

Trait distribution among H4 progeny. The green, red and black full dots represent the parental values of the strains B-1A, G-4A and H4.

### Additional File 3

Correlation analysis within each trait investigated in the H4 progeny. The test applied was the Pearson test. The size and the color of the dots represent the pvalue and the correlation rate, respectively. Only significant correlations corrected p values (BH) lower than 0.001 were shown.

### Additional File 4

SNP detected by NGS (sheet 1); Tiling (Sheet 2) and filtered markers with their occurrence in the 7 segregants genotyped (sheet 3)

### Additional File 5

KASP Markers used for linkage analysis

### Additional File 6

List on SNP within parental strains and their effect for loci on chromosomes VIII and IV.

### Additional File 7

Biomass viability for the hemyzygous hybrids Δ*OYE2^B^*::*KanMx4/OYE2^G^* (red) and *OYE2*^B^/Δ*OYE2^G^::KanMx4* (green) and H4 (black). The dots represent mean value for the sampling points and the shaded area the standard deviation estimated with at least five repetitions.

### Additional File 8

Panel A. The bar plots represent the average values of residual sugars after isothermal fermentations carried out at 24, and 32°C in the laboratory strain background (BY4741). The genotypes Δ*oye2* and *OYE2(wt)* were shown in blue and grey, respectively. Bars represent standard error of five repetitions, the statistical differences between the hemizygous was tested by a Wilcoxon-Mann-Whitney Test (the p value is coded as follow, ‘*’=p<0.05). Panel B. Fermentation kinetics (CO_2_ produced time course at 24°C.) for the same strains and with the same color key.

### Additional File 9

This figure illustrates the deletion observed in the strain B-1A for the genomic region encompassing the genes *FIT2, FIT3, FRE5, YOR385W, PHR1, YOR387C.* The deletion was found by comparing the read per kb observed for all the genes of the right arm of chromosome XV. Green bars (B-1A), red bars (G-4A).

### Additional File 10

#### Reliability of the enzymatic assay of glucose and fructose

Panel A. Linearity of the assay. The values shown are the average of two repetitions and the error bar indicates the standard deviation. The blue line indicates the linear regression line, the adjusted R-Squared is indicated. Panel B. Recovery of the assay. Each point represents the average value of three repetitions. The error bars indicate the standard deviation. The blue line indicates the linear regression line. Recovery is indicated.

Linearity of the assay (panel A). The concentration of a sample was measured at different dilution levels (1/100, 1/200 and 1/400). Linear regression with a R-Squared close to 1 indicates the linearity of the enzyme assay in this range (0.15 g/L −0.6g/L).

Recovery of the assay (panel B). Different concentrations of glucose or fructose are added to a sample (0.4 g/L, 1 g/L, 10 g/L and 15 g/L). The slope of the linear regression line indicates which part of the added concentration is actually measured (recovery). A slope close to 1 shows a good recovery of the assay between 0.4 g/l and 15 g/l.

### Additional File 11

Details of the gene deletion method

## Declaration section

### Data availability and materials

The datasets supporting the conclusions of this article are included within the article (6 tables, 6 figures) and its 11 additional files.

### Competing interests

MB, CM, EP and PM are employed by BIOLAFFORT company. This does not alter the authors’ adherence to all the journal policies on sharing data and materials.

### Consent for publication

Not applicable.

### Ethics approval and consent to participate

Not applicable.

### Funding

PM received 2 grants from Conseil Regional d’Aquitaine (SAGESSE for genome sequencing, and QTL2 for genotyping). The funders had no role in study design, data collection and analysis, decision to publish, or preparation of the manuscript.

### List of abbreviations

SNP: Single Nucleotide Polymorphism
PCD: Programmed Cell Death
ROS: Reactive Oxygen species
QTL: Quantitative Trait Loci
INDEL: Insertion DELetion
NIL: Nearly Isogenic Line
RHA: Reciprocal Hemizygous Assay

## Authors’ contributions

PM and DD conceived the work, PM, EP, PD, MB and CM realized experiments, PM wrote the manuscript.

## Acknowledgements

Not applicable.

