## Supplementary figures and images for "Natural allelic variations of *Saccharomyces cerevisiae* impact stuck fermentation due to the combined effect of ethanol and temperature; a QTL-mapping study"

### additional file 1

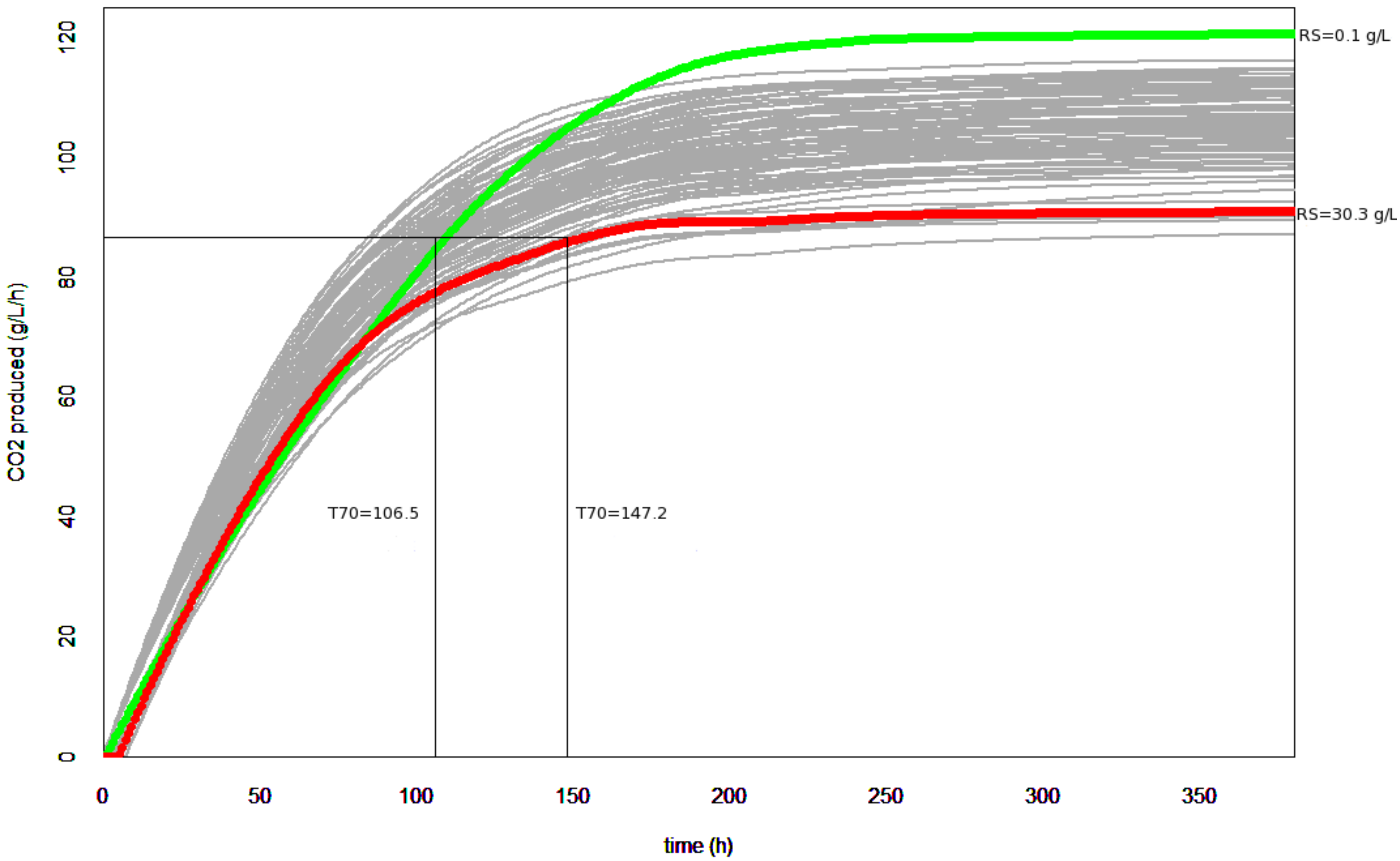

### additional file 2

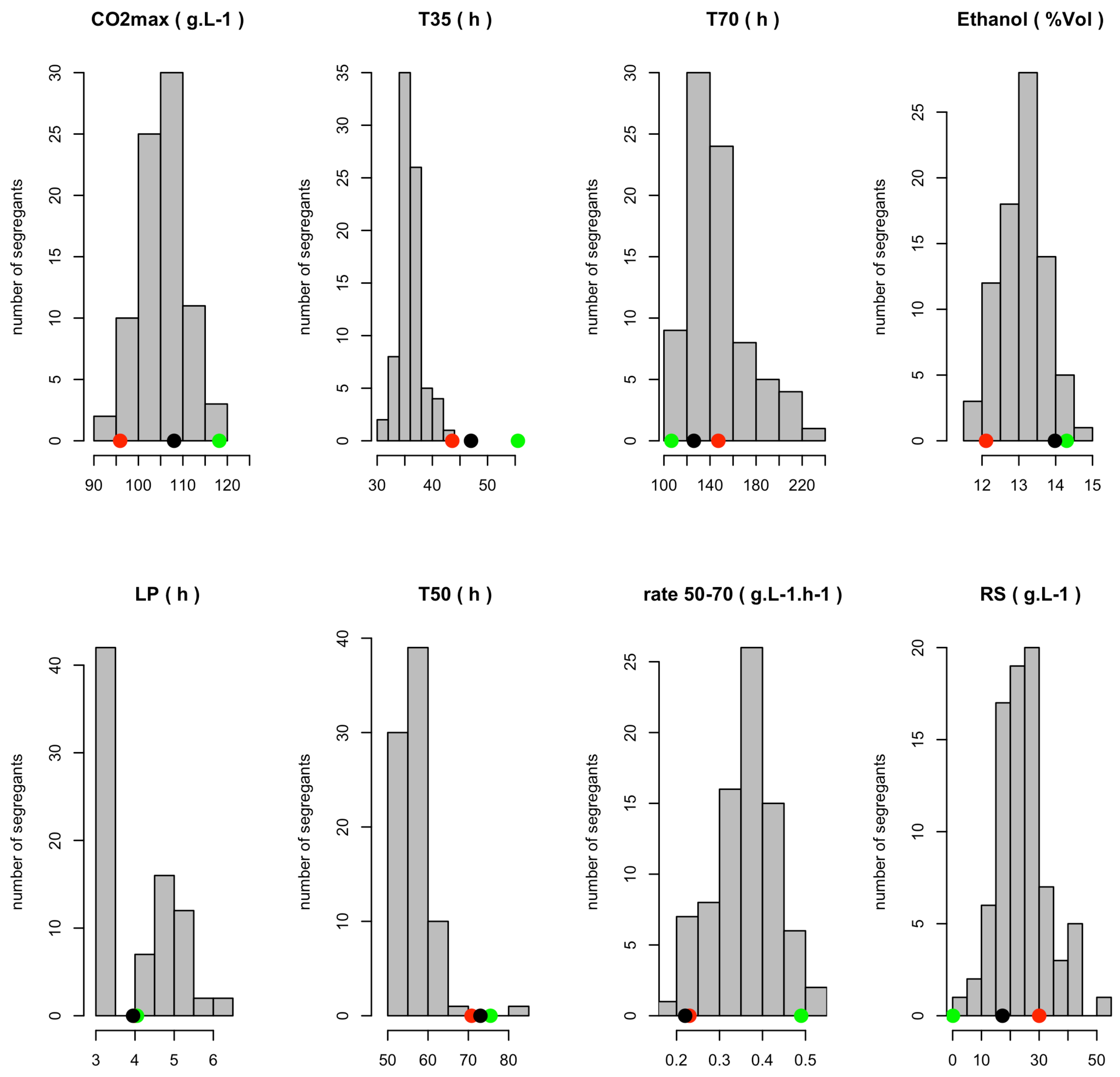

### additional file 3

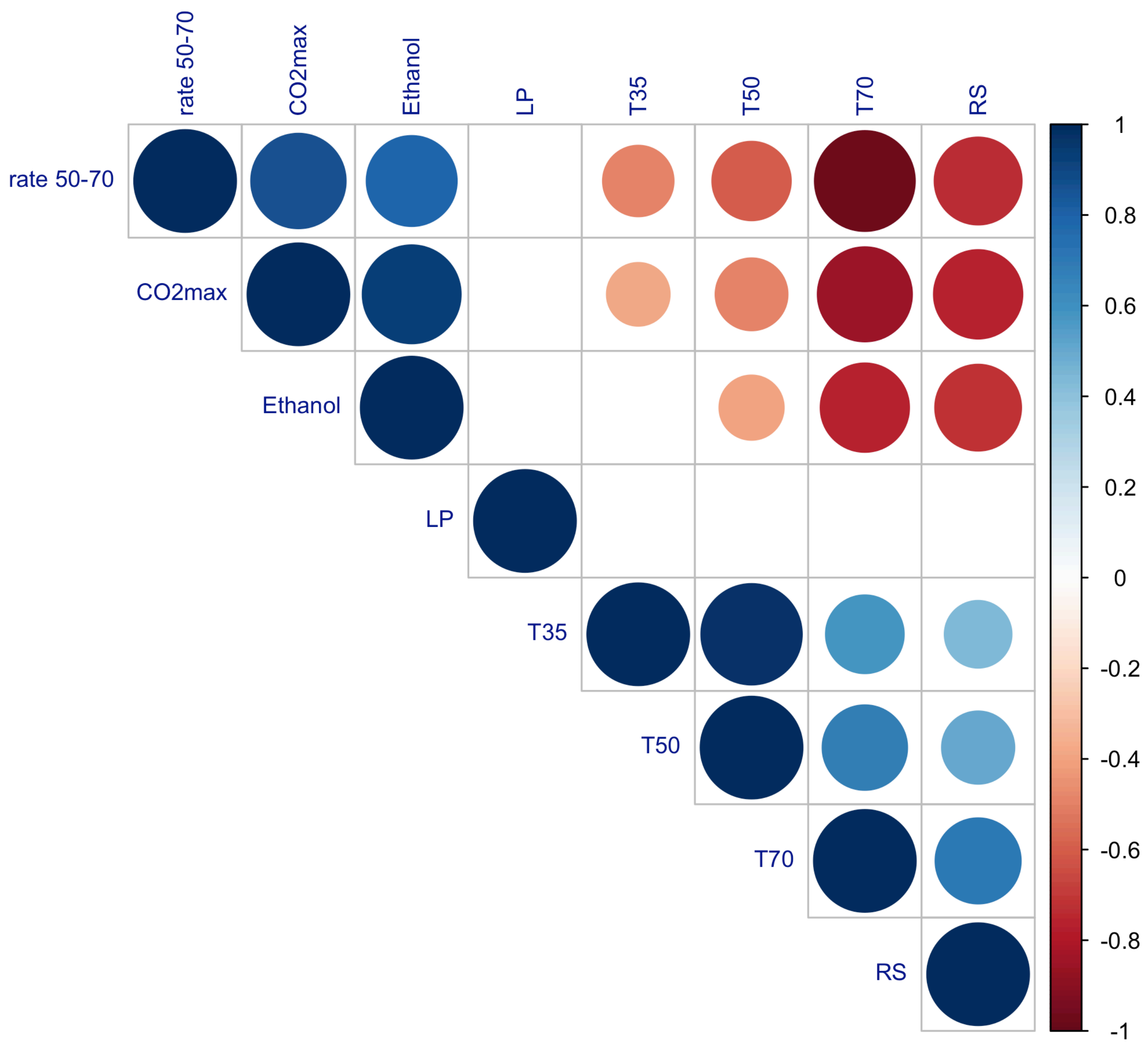

### additional file 7

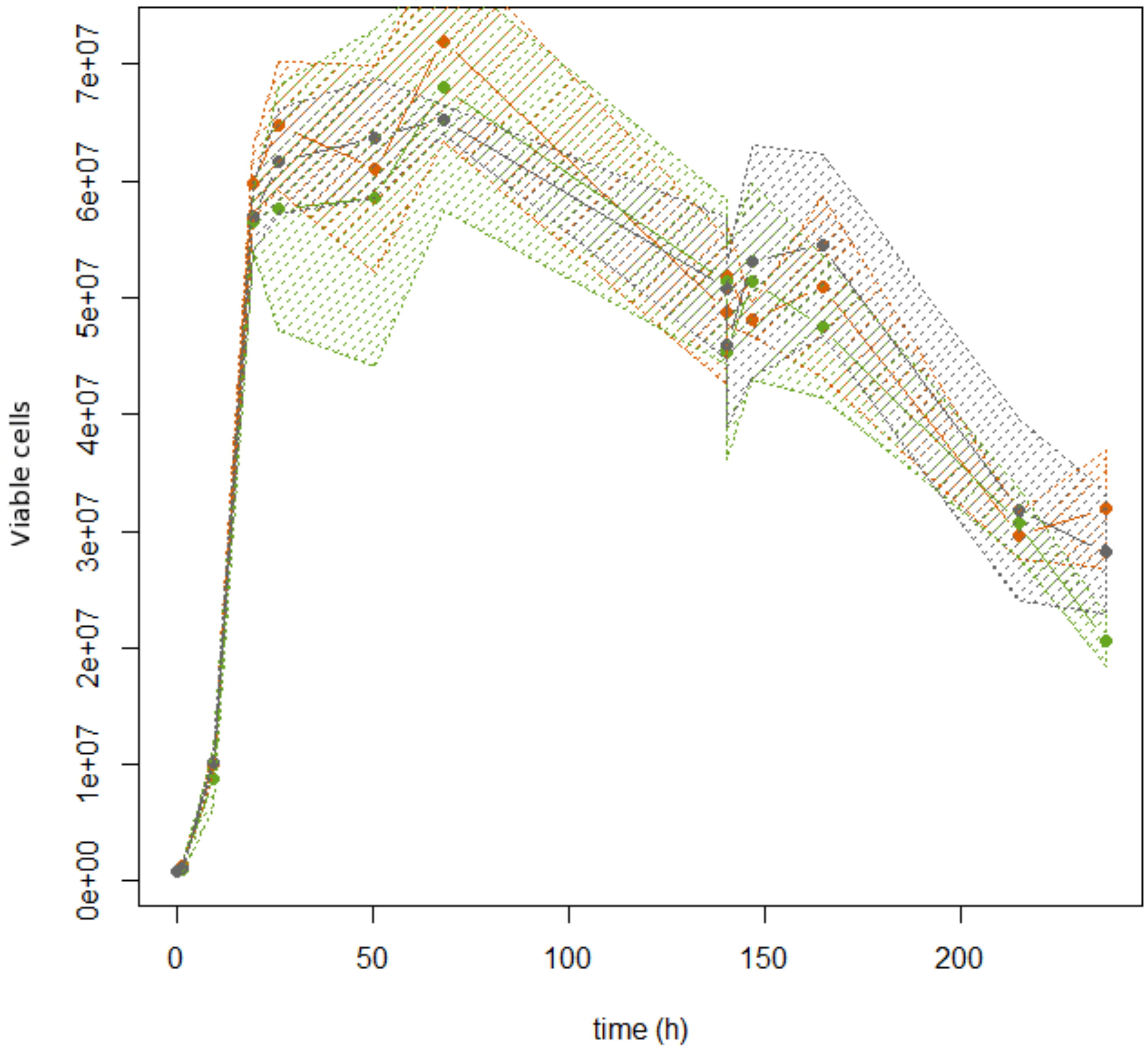

### additional file 8

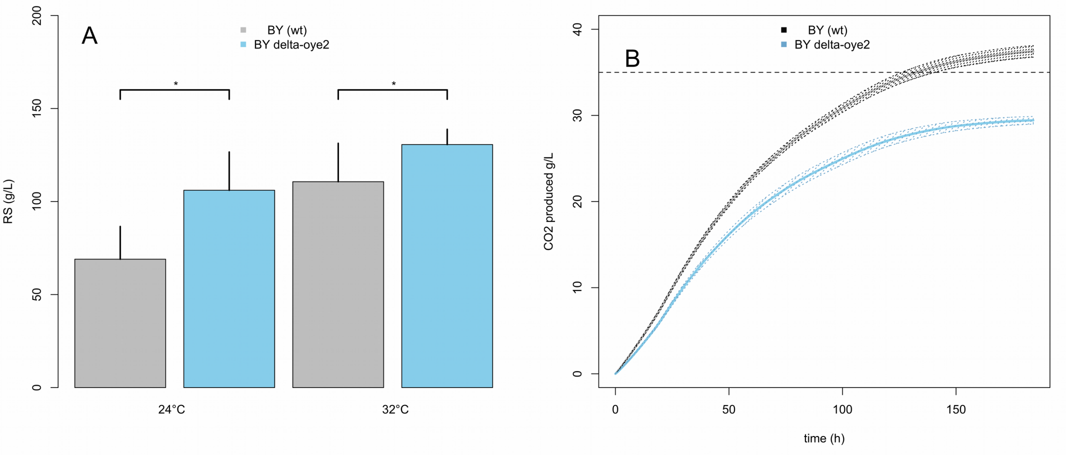

### additional file 9

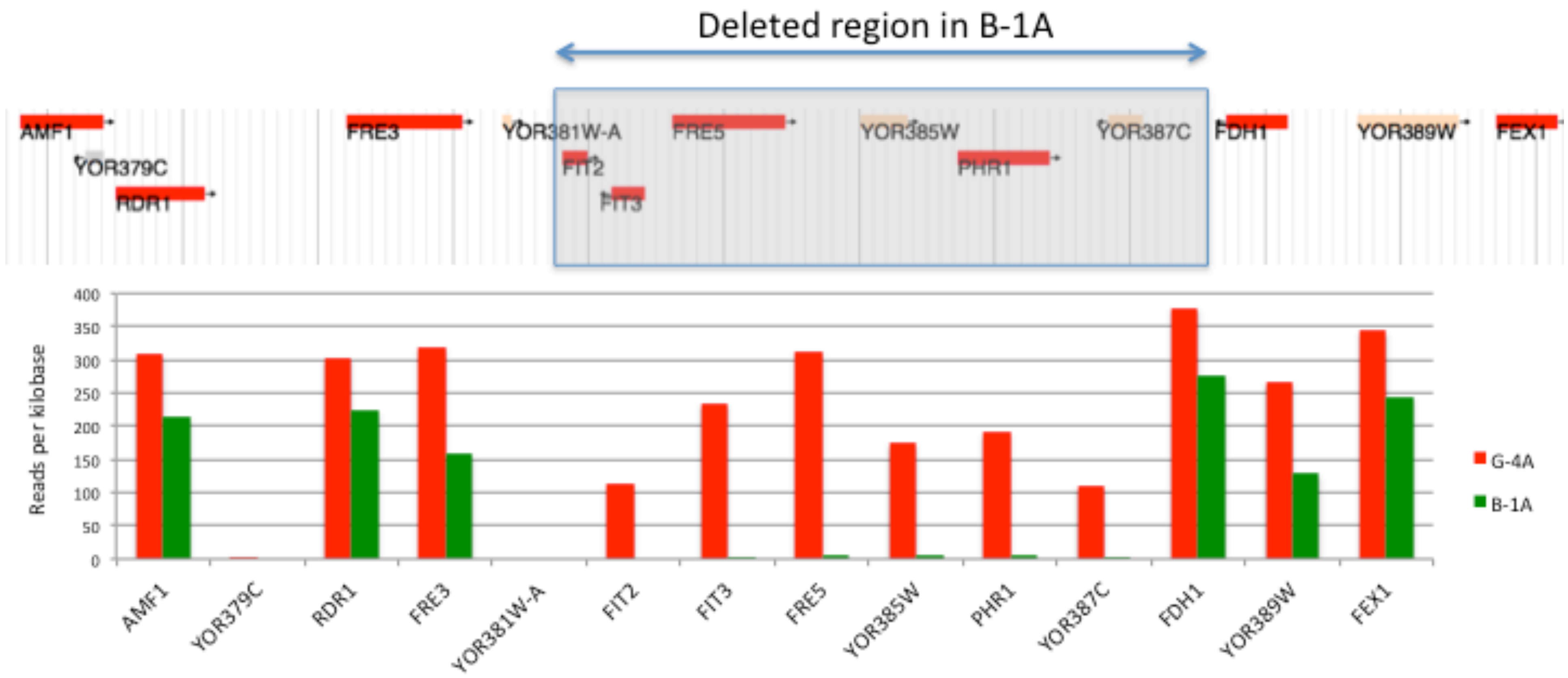

### additional file 10

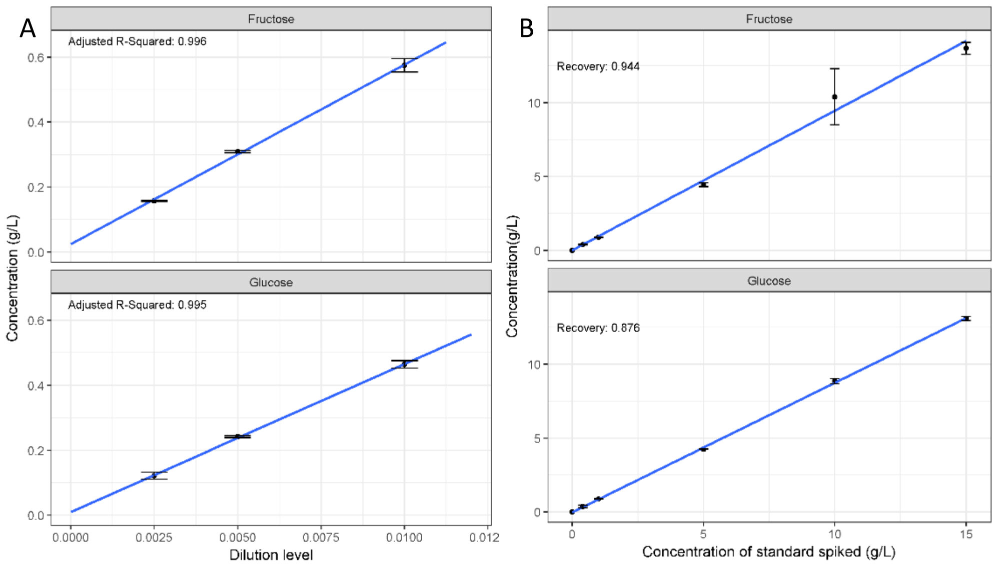
