## additional file 11 for "Natural allelic variations of *Saccharomyces cerevisiae* impact stuck fermentation due to the combined effect of ethanol and temperature; a QTL-mapping study"

### File S7 MOLECULAR METHODS DETAILS

#### 1. PCR TO AMPLIFY DELETION CASSETTES TO TRANSFORM

*KAN* cassette

≈500pb

≈500pb

- DNA extracted using Wizard genomic DNA purification kit (Promega).
- Primers used:

Table 1. Primers used to amplify each gene specific deletion cassette

| primer identification | sequence | Tm | Gene deletion cassette |
| --- | --- | --- | --- |
| P1035 | **AGTTAATTTGCCAAGCAGCG** | 55.3 | *VHS1_*fwd |
| P1036 | **TCAATTCAGGGATCTGGAGTT** | 55.9 | *VHS1_*rev |
| P918 | **CTTGGATGCTCATTACAAATGAA** | 57.1 | *OYE2_*fwd |
| P919 | **ATTACACGCGGTGTAAGAAGATG** | 59.4 | *OYE2_*rev |

- PCR mix:
  - 10 uL iProof buffer,
  - 1 µL dntp mix (10 mM)
  - 2.5 uL primer forward (20 µM),
  - 2.5 uL primer reverse (20 µM),
  - 31 µL uL sterile H_2_O,
  - 1 uL DNA template (50 – 500 mg).
  - 0.5 µL iProof Polymerase
- PCR program:

PCR conditions to amplify deletion cassettes:

- *VHS1*

| 98 °C 30 s | **35x** | 72 °c 5 min | 10 °c infinite |
| --- | --- | --- | --- |
|  | 98 °c 30 s |  |  |
|  | 57 °c 30 s |  |  |
|  | 72 °c 1 min 30 s |  |  |

- *OYE2*

| 98 °C 30 s | **35x** | 72 °c 5 min | 10 °c infinite |
| --- | --- | --- | --- |
|  | 98 °c 30 s |  |  |
|  | 58 °c 30 s |  |  |
|  | 72 °c 1 min 30 s |  |  |

- Verification correct amplification on agarose gel 1%

#### 3. PCR TO VERIFY TRANSFORMATION

Before transformation:

Gene X

p forward

≈600pb

After transformation

*KAN* cassette

p reverse:

**p560 bis**

p forward

≈600pb

The PCR verification consists in using one primer at 600pb (approx.) from the deleted gene loci and one primer that will anneal inside the *KAN* cassette. Thus, we verify the correct insertion of the KAN cassette in the right loci.

- PCR mix:
  - 4ul Taq&GO,
  - 0.1 primer forward (20 µM),
  - 0.1 primer reverse (20 µM)
  - 1 µL DNA template ((50 – 500 mg)
  - 14.8 H2O.
- PCR program (*VHS1*):

| 95 °c 10 min | **35x** | 72 °c 5 min | 10 °c infinite |
| --- | --- | --- | --- |
|  | 95 °c 45 s |  |  |
|  | 54 °c 45 s |  |  |
|  | 72 °c 2 min |  |  |

- PCR program (*OYE2*):

| 95 °c 10 min | **35x** | 72 °c 5 min | 10 °c infinite |
| --- | --- | --- | --- |
|  | 95 °c 45 s |  |  |
|  | 57 °c 45 s |  |  |
|  | 72 °c 2 min |  |  |

- Primers used:

Table 2. Primers used in the verification of the transformation.

| Primer id | sequence | Tm | Gene |
| --- | --- | --- | --- |
| P560bis | CGGCGCAGGAACACTG | 56.9 | *KAN* test insertion  Anneals at the middle of the *KAN* cassette. |
| P1042 | **GTGTATCTAGGTCCGCG** | 55.2 | *VHS1* test insertion |
| P920 | **GCGGCATGCTTTTTCCGT** | 58.8 | *STR2* test insertion |

- Verification in agarose gel 1%.

#### 4. ALLELE CONTROL

Methods used for checking the remaining allele after KO

*VHS1 sequencing the fragment generated by the following PCR*

| P1033 | ACAAGCTTCACCAATGAAGGC | 55.3 | SNP CHECK foward |
| --- | --- | --- | --- |
| P1034 | **TGTGTCTAGCGTTCCGAAACT** | 55.9 | SNP CHECK foward |

1) PCR to amplifly fragment to be sequenced

PCR mix:

4 uL Taq&GO,

0.1 primer forward (20 µM),

0.1 primer reverse (20 µM),

1 uL genomic DNA

14.8 H2O.

Program:

| 95 °c 10 min | **35x** | 72 °c 5 min | 10 °c infinite |
| --- | --- | --- | --- |
|  | 95 °c 45 s |  |  |
|  | 54 °c 45 s |  |  |
|  | 72 °c 1 min |  |  |

*OYE2 was typed by PCR using the following primers the forward primer is common while the reverse are specific to the mutation and lead to a clear cut presence/absence of amplification*

| 95 °c 5 min | **35x** | 72 °c 10 min | 10 °c infinite |
| --- | --- | --- | --- |
|  | 95 °c 43 s |  |  |
|  | 65 °c 35 s |  |  |
|  | 72 °c 45 min |  |  |

| P794 | **GCTGACATTTTGTAGAAAGTGTCTCTGTC** | 63,9 | OYE2 Forward |
| --- | --- | --- | --- |
| P795 | **ttgaacctcgtgtcaccaacccattt** | 63.2 | OYE2 B1 specific reverse |
| P796 | **ttgaacctcgtgtcaccaacccatta** | 63.2 | OYE2 G4 specific reverse |

1) PCR to amplifly fragment to be sequenced

PCR mix:

4 uL Taq&GO,

0.1 primer forward (20 µM),

0.1 primer reverse (20 µM),

1 uL genomic DNA

14.8 H2O.

Program:

Verification in agarose gel 2%.
